# An enterococcal phage protein broadly inhibits type IV restriction enzymes involved in antiphage defense

**DOI:** 10.1101/2023.11.16.567456

**Authors:** Nathan P. Bullen, Cydney N. Johnson, Shelby E. Andersen, Garima Arya, Sonia R. Marotta, Yan-Jiun Lee, Peter R. Weigele, John C. Whitney, Breck A. Duerkop

## Abstract

The prevalence of multidrug resistant (MDR) bacterial infections continues to rise as the development of antibiotics needed to combat these infections remains stagnant. MDR enterococci are a major contributor to this crisis. A potential therapeutic approach for combating MDR enterococci is bacteriophage (phage) therapy, which uses lytic viruses to infect and kill pathogenic bacteria. While phages that lyse some strains of MDR enterococci have been identified, other strains display high levels of resistance and the mechanisms underlying this resistance are poorly defined. Here, we use a CRISPR interference (CRISPRi) screen to identify a genetic locus found on a mobilizable plasmid from *Enterococcus faecalis* involved in phage resistance. This locus encodes a putative serine recombinase followed by a Type IV restriction enzyme (TIV-RE) that we show restricts the replication of phage phi47 in *E. faecalis*. We further find that phi47 evolves to overcome restriction by acquiring a missense mutation in a TIV-RE inhibitor protein. We show that this inhibitor, termed type IV restriction inhibiting factor A (*tifA*), binds and inactivates diverse TIV-REs. Overall, our findings advance our understanding of phage defense in drug-resistant *E. faecalis* and provide mechanistic insight into how phages evolve to overcome antiphage defense systems.

## INTRODUCTION

Typically members of the gut microbiota of humans and other animals, the enterococci are Gram-positive bacteria that can cause life-threatening disease following antibiotic-mediated dysbiosis^1,2^. During antibiotic treatment, multidrug-resistant (MDR) *Enterococcus faecalis* can outgrow antibiotic-sensitive members of the microbiota and this increases the likelihood that MDR *E. faecalis* will breach the intestinal barrier and enter the bloodstream leading to infection^1,2^. These MDR infections are difficult to treat due to limited antibiotic options and consequently, bacteriophages or phages (viruses that infect bacteria) are increasingly being considered as a therapeutic option as the frequency of MDR infections continues to rise^3–5^. Phage therapy offers advantages over traditional antibiotic therapy because phages exhibit strain-level specificity, which reduces off-target bacterial killing, they are self-limiting following the exhaustion of their bacterial host, and their abundance in nature implies that a phage specific to a target bacterium can usually be identified^6–10^.

As is the case with the emergence of antimicrobial resistance, exposure of bacteria to phages also imposes a strong selective pressure that leads to the evolution or acquisition of mechanisms to resist infection. Recent studies have shown that bacteria have evolved diverse antiphage defense systems to prevent successful phage infection^11–13^. These systems are often concentrated in bacterial genomes within so-called “defense-islands”, which are characterized by the clustering of genes encoding two or more defense systems and are typically carried on mobile genetic elements, including integrative and conjugative elements, plasmids, transposons, and temperate phages^14–16^. This capacity to mobilize and spread between bacterial species represents a robust pan-genomic immune system that contributes to the rapid evolution of resistance to phage infection and constitutes a significant challenge to the effective application of phage therapy.

Antiphage defenses utilize a variety of mechanisms to overcome phage infection. These include restriction-modification systems in which host proteins epigenetically label nucleobases and/or the phosphate backbone to distinguish host DNA from phage DNA, RNA-based adaptive nucleic acid targeting systems such as CRISPR-Cas^16–21^, abortive infection systems and other phage interference systems that stop phage replication and spread^22–26^, chemical defenses that restrict phage infection^27^, and phage receptor alterations that abrogate the ability of phages to interact with the bacterial cell surface^28–32^. In turn, phages have evolved counter-defenses to antiphage defense systems including enzymes that modify tail proteins for bacterial cell recognition, DNA modification to avoid restriction systems, and phage-encoded proteins that interfere with antiphage defenses^33–37^. Compared to bacterial antiphage defense systems, the breadth of phage-encoded proteins that support counter-defenses and the mechanisms that drive their evolution are poorly understood. Thus, prior to phages being widely used as therapeutics for MDR bacterial infections, a detailed understanding of the mechanisms by which bacteria subvert phage infection and, conversely, how phages overcome these mechanisms is needed.

In this study, we sought to understand why some MDR strains of *E. faecalis* are susceptible to infection by virulent enterococcal phages whereas others exhibit resistance^38^. By conducting a CRISPR interference (CRISPRi) screen on one such resistant strain, we pinpoint a mechanism of resistance to a two-gene locus residing on a <u>p</u>heromone responsive <u>p</u>lasmid (PRP) that inhibits infection of the enterococcal phage phi47. Further analysis revealed that resistance to phi47 infection is solely conferred by the activity of one of these genes, EF_B0059, encoding a putative type IV restriction enzyme (TIV-RE). We subsequently isolated phages that are resistant to restriction by EF_B0059 and discovered that the *orf65* gene of phi47 encodes a small protein that can evolve to not only bind and inactivate EF_B0059, but also other highly divergent type IV restriction enzymes. From this, we posit that *orf65* encodes a rapidly evolving and broadly acting anti-TIV-RE protein. Overall, our data demonstrate that *E. faecalis* utilizes a mobilizable antiphage system to resist phage infection, and that phages escape this restriction through a coevolving small open reading frame found in enterococcal and related phages.

## RESULTS

### The pheromone responsive plasmid pTEF2 is sufficient to inhibit phage infection

In our attempts to understand the mechanisms underlying phage-host interactions in MDR enterococci, we previously observed that the lytic phage phi47 can infect *E. faecalis* strain SF28073 but not the closely related strain V583^38^. phi47 readily adsorbs to both SF28073 and V583, indicating that this resistance cannot be attributed to differences in the cell surface of these strains preventing phage binding^38^. From this, we hypothesized that *E. faecalis* V583 may encode one or more antiphage defense systems that restrict the replication of phi47.

A comparison of the *E. faecalis* SF28073 and V583 genomes revealed that they are very similar (nearly 90% sequence identity across shared sequences) (Fig. S1A). However, sequence comparison of the three natural plasmids carried by these strains showed that the plasmids of V583 (pTEF1, pTEF2, pTEF3) differ significantly from that of SF28073 (pSF1, pSF2, pSF3) (Fig S1B). Based on these observations, we hypothesized that one or more plasmid(s) of V583 may be responsible for resistance to phi47 infection. To explore this idea, we infected *E. faecalis* V19, a plasmid-cured derivative of V583, with phi47^39,40^. Unlike V583, V19 supports robust replication of phi47 indicating that one or more of the three V583 plasmids confers resistance to phi47 (Fig. 1A). Of these three plasmids, pTEF1 and pTEF2 are PRPs that mobilize through conjugation upon sensing a peptide pheromone signal produced by plasmid-free cells^41^. In contrast to pTEF1 and pTEF2, pTEF3 is a comparatively small, non-mobilizable plasmid^42^. Given that plasmid mobilization has been previously demonstrated to disseminate antiphage defenses^43–45^, we began our investigation with pTEF1 and pTEF2.

**Figure 1.**
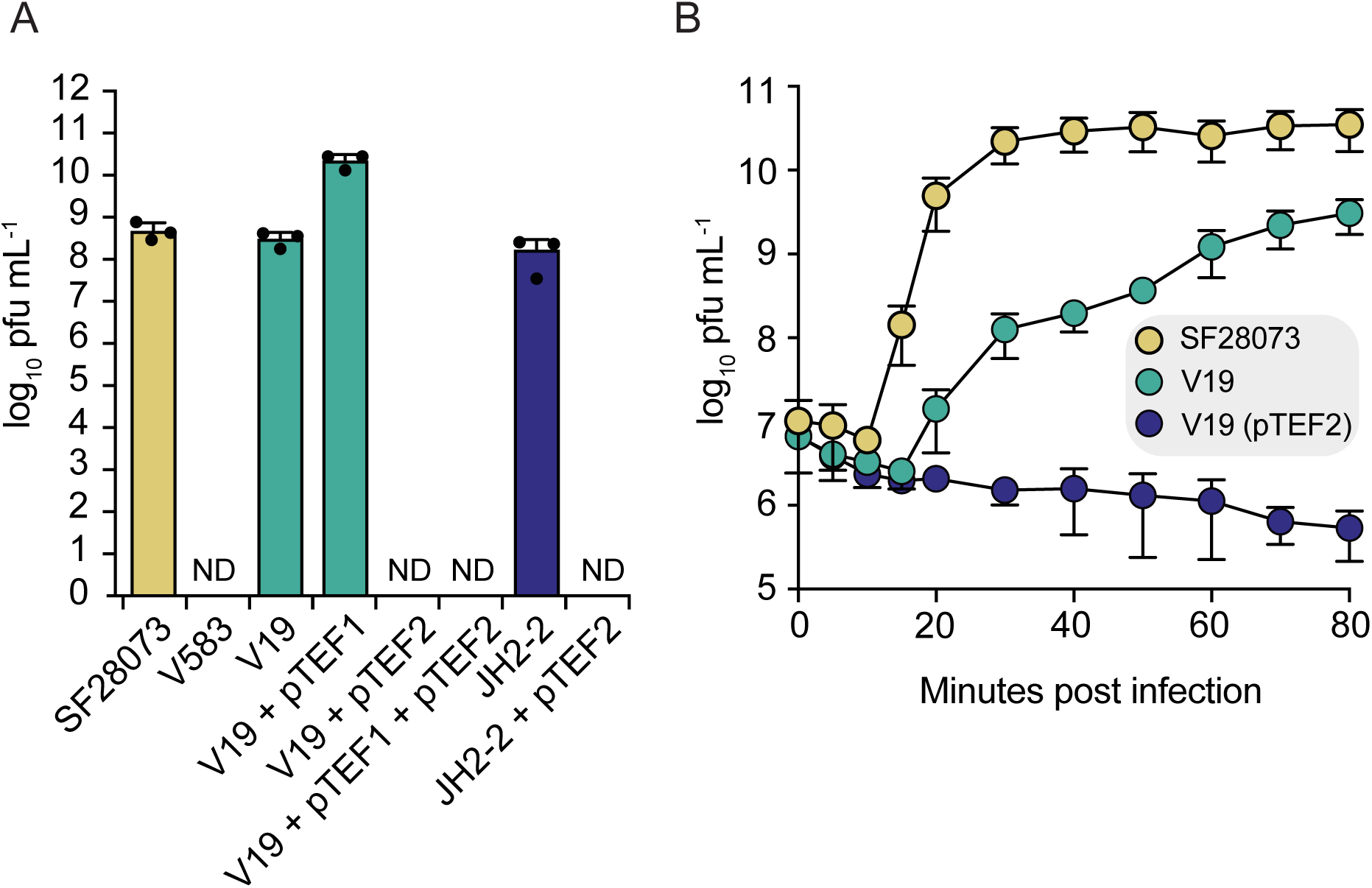
The pheromone responsive plasmid pTEF2 is sufficient to inhibit phi47 replication. **(A)** Capacity of phage phi47 to form plaques measured by log_10_ PFU/mL on the indicated *E. faecalis* strains. Data represent mean ± s.d. of three technical replicates. **(B)** PFU/mL produced upon phi47 infection of indicated *E. faecalis* strains over 80 minutes. All strains were infected with an equivalent MOI of 1.0. Data represent mean ± s.d. of four technical replicates. Source data are provided as a Source Data file.

To determine if an *E. faecalis* V583 mobilizable plasmid mediates resistance to phi47, we conjugated pTEF1 and pTEF2 into the phi47 susceptible *E. faecalis* V19 strain. Subsequent infections with phi47 showed pTEF1 had no effect on the ability of phi47 to replicate in V19 whereas the carriage of pTEF2 completely restricted phi47 replication (Fig. 1A). This pTEF2-mediated resistance to phi47 is independent of the *E. faecalis* strain background because mobilization of pTEF2 into *E. faecalis* JH2-2, which shares 87% of its genome at a nucleotide identity of 99% or greater with V583^46^, was sufficient to prevent replication of phi47 (Fig. 1A). To confirm that no infectious phages were being released from pTEF2-containing cells, we also measured total infectious phi47 particle production during infection (Fig. 1B). While the strains V19 and SF28073 support the production of infectious phi47 particles, V19 pTEF2 grown in the presence of phi47 did not show an increase in recoverable phages over time. This data demonstrates that pTEF2 alone confers resistance to phi47.

Some antiphage defense systems are predicted to function through a mechanism termed abortive infection whereby the activation of the antiphage system stops the spread of virus at the expense of the viability of the bacterial host^22,23,47–49^. Such population-level resistance can be measured by infecting bacteria with phage at high <u>m</u>ultiplicities <u>o</u>f infection (MOI) because this results in the complete collapse of the bacterial culture. In contrast, some defense systems confer individual-level resistance to phage, often by directly targeting phage replication processes during early infection. In these instances, bacterial growth upon phage infection is comparable at low and high MOIs. To determine if phi47 resistance is due to abortive infection, we measured the optical density (OD_600_) of *E. faecalis* cultures during infection by phi47 at an MOI of 10 (Fig. S2A). In doing so, we observed that V19 pTEF2 cells continued to grow during phi47 infection at a similar rate to that of uninfected V19 pTEF2. Thus, pTEF2 antiphage defense is not mediated by the death or inability of the bacteria to grow, but instead provides direct immunity to phi47. Additionally, we confirmed that phi47 adsorbs equally to the cell surface of both V19 and V19 pTEF2 compared to V583 (Fig. S2B). Taken together, our data suggests that the pheromone responsive plasmid pTEF2 encodes an antiphage defense system that prevents phage replication in actively growing cells.

### A GmrSD-like protein encoded on pTEF2 is responsible for phage restriction

We next determined which gene(s) on pTEF2 were responsible for the restriction of phi47. Of the 62 open reading frames (ORFs) encoded by pTEF2, approximately half are predicted to be involved in the maintenance and conjugation of the plasmid. Furthermore, more than half of the genes on pTEF2 share 95-97% sequence identity to another *E. faecalis* PRP, pCF10, which, unlike pTEF2, does not confer resistance to phi47 when conjugated into V19 (Fig. 2A and 2B). Comparing the sequences of pTEF2 and pCF10 revealed that pTEF2 encodes a unique 15 kb region encompassing 14 ORFs not present in pCF10 (EF_B0048 to EF_B0065). We reasoned that the gene(s) involved in phi47 restriction were therefore likely encoded by one or more these 14 ORFs.

**Figure 2.**
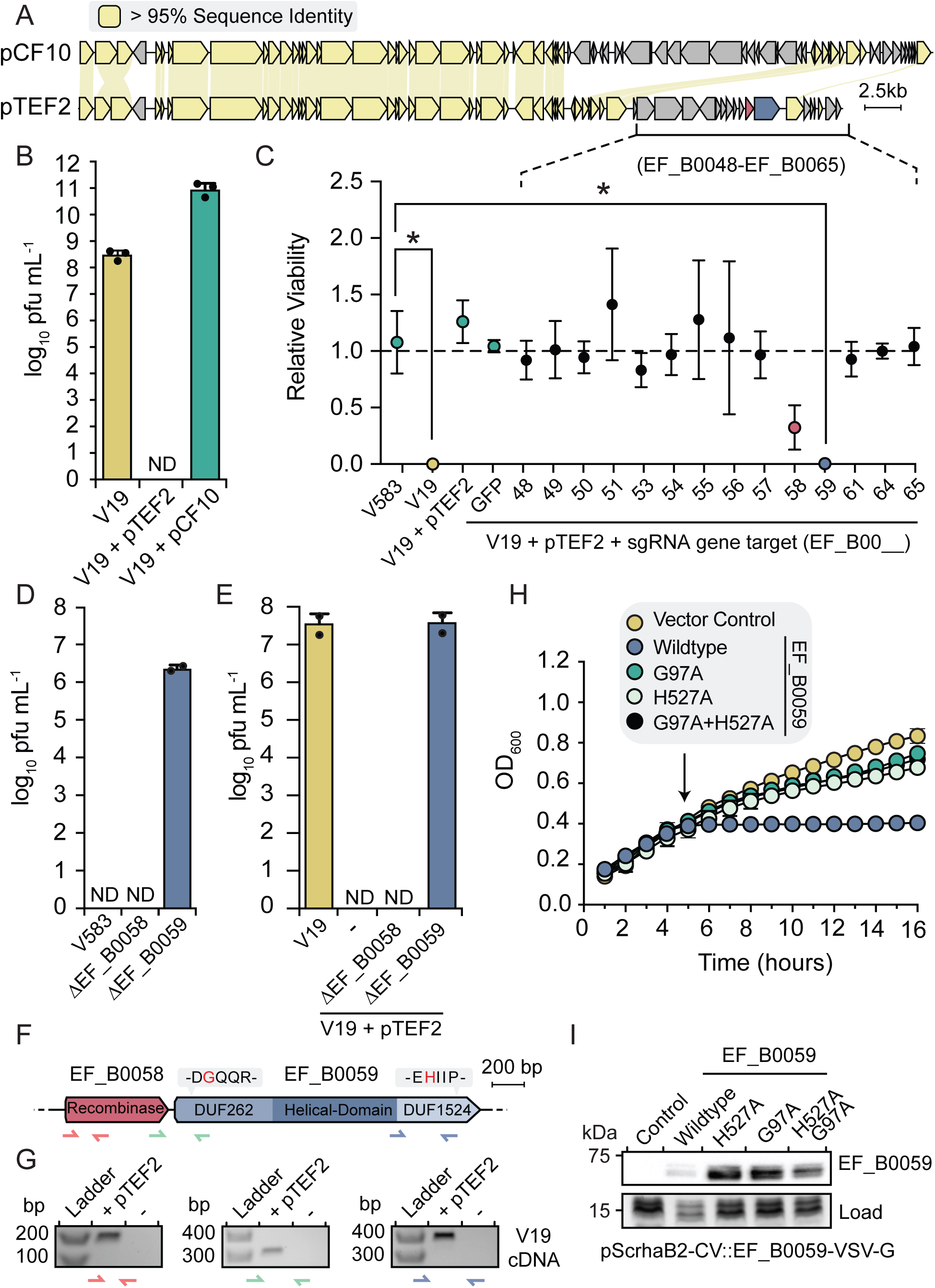
*EF_B0059* encodes a putative type IV restriction enzyme that prevents infection by phi47. **(A)** Sequence alignments between the pheromone responsive plasmids pTEF2 and pCF10. Genes highlighted in yellow indicate a sequence identify greater than 95%. **(B)** Capacity of phage phi47 to form plaques measured by PFU/mL on the indicated *E. faecalis* strains. Data represent mean ± s.d. of three technical replicates. **(C)** Relative viability of *E. faecalis* V19 pTEF2 cells upon transcription knock-down of the indicated pTEF2 encoded genes of interested using CRISPR interference. Relative viability was determined by comparing the growth of each strain in the presence and absence of phi47. Statistical significance determined by ANOVA with multiple corrections, *p=0.0004. Data represent mean ± s.d. of four technical replicates. **(D)** and **(E)** Outcome of phi47 infections measured by PFU/mL in *E. faecalis* strains V583 (D) and V19 (E) in which the genes encoding EF_B0058 or EF_B0059 on pTEF2 are deleted. Data represent mean ± s.d. of two technical replicates. **(F)** Schematic of the EF_B0058 and EF_B0059 genes. **(G)** Reverse transcription-PCR analysis of the EF_B0058 and EF_B0059 mRNA transcript in *E. faecalis* V19 with or without pTEF2. Primer sets used to probe the structure of the EF_B0058-EF_B0059 transcriptional operon are shown in red (EF_B0058 specific), green (intergenic) and blue (EF_B0059 specific). **(H)** Growth curves of *E. coli* expressing EF_B0059, the indicated active site mutants, or empty vector. Arrow indicates when protein expression was induced. Data represent mean ± s.d. of three technical replicates. **(I)** Western blot of VSV-G-tagged EF_B0059 in *E. coli* strains from (H), 30 minutes post-induction. A non-specific band from the PAGE of these samples stained with Coomassie Brilliant Blue is used as a loading control (load). Data are representative of three independent experiments. Source data are provided as a Source Data file.

To test this prediction, we performed a CRISPR interference (CRISPRi) gene knockdown screen in *E. faecalis* V19 pTEF2. CRISPRi uses a catalytically inactive Cas9 (dCas9) that binds to target DNA sequences when directed by complementary small guide RNAs (sgRNAs), resulting in transcriptional repression of target genes^50^. We used plasmids expressing dCas9 and sgRNAs that targeted each of the 14 ORFs unique to pTEF2 and transformed these into V19 pTEF2 cells to selectively knock down target gene expression. The relative viability of the resulting transformants was then measured in the presence or absence of phi47 (Fig. 2C). Strikingly, under these experimental conditions, the sgRNA-dependent repression of either EF_B0058 or EF_B0059 in the presence of phi47 resulted in a significant decrease in cell viability when compared to V19 pTEF2. V19 transformed with an sgRNA targeting *gfp* (sgRNA-GFP) was used as a non-specific control. To validate the findings of our screen, we generated single gene knock outs of either EF_B0058 or EF_B0059 in both V583 and V19 carrying pTEF2 (Fig. 2D and 2E). Subsequent infections with phi47 in these strains revealed that the deletion of EF_B0058 did not restore the ability of phi47 to replicate in V583 or V19 whereas deletion of EF_B0059 did. This result indicates that EF_B0059 alone confers resistance to phi47 infection. A closer examination of the genetic context of EF_B0058 and EF_B0059 found that these genes are encoded in tandem and are separated by a small intergenic region (54 base pairs), suggesting that they may be encoded on the same mRNA (Fig. 2F). To test this prediction, we performed reverse-transcription PCR to generate cDNA from total RNA purified from *E. faecalis* V19 with and without pTEF2. PCR reactions using this cDNA as a template and primers that span the intergenic region between EF_B0059 and EF_B0058 revealed that EF_B0058 and EF_B0059 exist on the same mRNA (Fig. 2G). From this finding, we concluded that the sgRNA targeting EF_B0058 likely also disrupts the expression of downstream EF_B0059, resulting in a reduced capacity to restrict phi47.

Bioinformatic analyses, including the generation of a high-confidence Alphafold2 model of EF_B0059, predict that this antiphage protein belongs to the GmrSD-like type IV modification-dependent restriction enzyme family (Fig. S3A and S3B). GmrSD family proteins are defined by N-terminal DUF262 and C-terminal DUF1524 domains connected by a small α-helical domain of variable length (Fig. S3A)^51^. DUF262 domains function as NTPases and contain a conserved (I/V)DGQQR motif that forms a nucleotide binding pocket whereas DUF1524 domains are structurally similar to HNH endonucleases and contain a conserved EHxxP catalytic motif found in members of the broad His-Me finger endonuclease superfamily^51,52^. EF_B0059 possesses an intact DUF262 VDGQQR motif but has a glutamate in place of the conserved proline in its DUF1524 EHxxP motif (Fig. S3A). The precise molecular details describing how these two domains coordinate DNA substrate recognition and cleavage are not yet understood, however, NTP hydrolysis by the N-terminal DUF262 has been shown to drive conformational changes that activate DUF1524’s nuclease activity resulting in DNA cleavage^51,53^. Moreover, these enzymes have been shown to target diverse types of modified DNA, in particular DNA containing modified cytosine bases including 5-methylcytosine^54,55^. Given this precedent, we expressed EF_B0059 in *E. coli* XL1-Blue, a K-12 derivative strain that contains 5-methylcytosine and N6-methyladenosine, under the assumption that overexpression of this putative TIV-RE would target the DNA of an evolutionarily distant bacterial species. In line with our hypothesis, we found that the heterologous expression of EF_B0059 is toxic to *E. coli* XL1-Blue (Fig. 2H and 2I). To test our assumption that EF_B0059 recognizes modified DNA for restriction, we repeated these intoxication experiments using the *E. coli* strain ER2796, which lacks the three active methyltransferases typically found in K-12 derivative strains, namely the Dam, Dcm, and *Eco*KI methylases. Surprisingly, we found that this strain still succumbs to intoxication by EF_B0059 (Fig. S4A). These data, combined with sequence analyses suggesting that the phi47 genome does not encode DNA modifying enzymes, indicate that EF_B0059 may not require modified phage DNA for restriction. To explore this possibility, we next examined the chemical structure of phi47 DNA. To this end, we isolated and digested DNA from phi47 virions and analyzed the resultant nucleotides by high-performance liquid chromatography coupled with mass spectrometry (Fig. S4B). Comparison of the phi47 DNA digestion products to that of nucleotide standards revealed that phi47 DNA is comprised of unmodified nucleotides. We also considered the possibility that EF_B0059 may act on phage DNA modifications conferred by the host cell from which the phage is propagated. To test this, we infected *E. faecalis* strains SF28073 and JH2-2 with phi47 and then measured the ability of the output phages to replicate in *E. faecalis* JH2-2 *+* pTEF2 (Fig. S4C). Regardless of the host strain used to propagate phi47, EF_B0059 prevented phage replication. When taken together with our biochemical data, this observation strongly suggests that phi47 DNA does not modify its DNA and that restriction of phage replication by EF_B0059 occurs through a modification-independent mechanism. Importantly, we found that EF_B0059 function is dependent on both its NTPase and nuclease activities because inactivating mutations to the conserved catalytic sites of either domain were non-toxic to *E. coli* XL1-Blue (Fig. 2H and 2I). Thus, EF_B0059 restriction presumably arises through its enzymatic activity acting on the phage genome. However, the mechanism by which EF_B0059 inhibits phi47 replication and how it discriminates between phage and host DNA remain to be determined.

### pTEF2-mediated resistance is overcome by a phage-encoded protein

Because phages are entirely dependent on their host for replication, they are under strong selective pressure to evolve ways to evade host defense systems. Thus, we next asked if phi47 can evolve to overcome EF_B0059-dependent restriction. To determine this, three phi47 single-plaque isolates were independently raised and purified to a titer of ~10^12^ PFU/mL. 1×10^9^ PFU of each isolate was plated on *E. faecalis* V19 pTEF2 and two resistant plaques were isolated for each replicate. Subsequent infections with the phages isolated from each of these plaques confirmed that they replicate equally on V19 and V19 pTEF2 (Fig. S5A). Intriguingly, whole genome sequencing of these phages found that they all harbor a single point mutation in the small open reading frame, *orf65*, that results in a nonsynonymous mutation at amino acid 32 from leucine to phenylalanine (Fig. 3A, Table S1). Mutations in phage genomes can arise to prevent DNA targeting defense systems from recognizing phage DNA substrate^56,57^. However, it is unlikely that any single change in DNA sequence would confer resistance given that restriction enzymes frequently target multiple sites within a phage genome^54,55^. We reasoned that the gene product of the mutated *orf65* may instead be responsible for evading EF_B0059 activity. To test this hypothesis, we infected *E. faecalis* V19 pTEF2 cells ectopically expressing either the wildtype or mutated Orf65 (Orf65^L32F^) with phi47 (Fig. 3B). These results confirmed that Orf65^L32F^, but not the wild-type protein, confers resistance to EF_B0059-dependent restriction.

**Figure 3.**
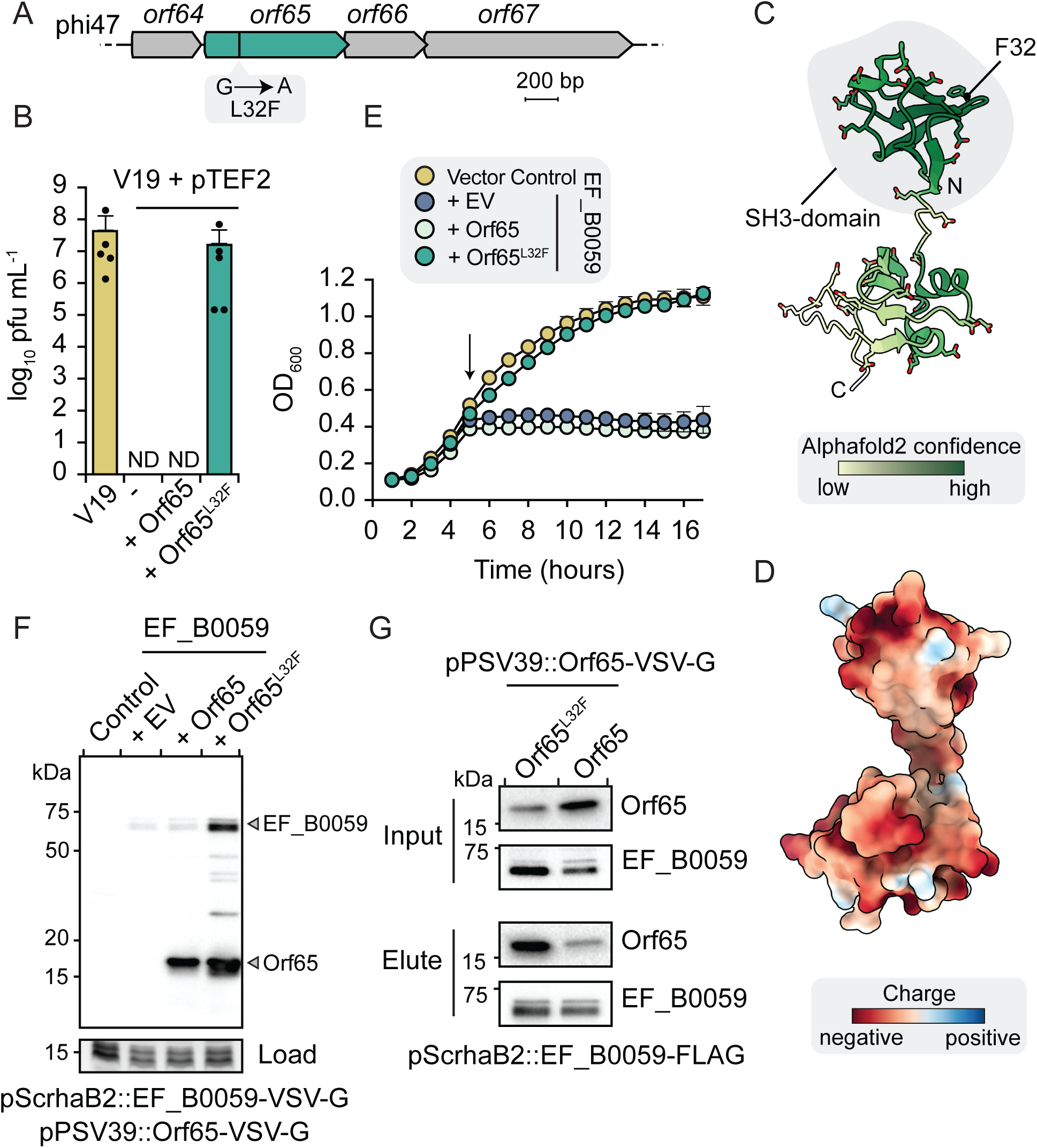
A mutation in phage phi47 protein Orf65 prevents EF_B0059-dependent restriction of phi47. **(A)** Gene schematic of the genomic loci of *orf65* in the phi47 genome. **(B)** Capacity of phi47 to form plaques measured by PFU/mL on *E. faecalis* strain V19 or V19 carrying pTEF2 and co-expressing Orf65 or Orf65^L32F^. Data represent mean ± s.d. of five technical replicates. **(C)** Ribbon representation of the predicted structure of Orf65 as generated using Alphafold2. The structure is colored according to the predicted local distance difference test (pLDDT). The distribution of the acidic residues throughout the structure is also shown. **(D)** The predicted amino acid surface electrostatic potential of Orf65. **(E)** Growth curves of *E. coli* expressing EF_B0059 and either Orf65 or Orf65^L32F^, or empty vector. The arrow indicates when protein expression was induced. Data represent mean ± s.d. of three technical replicates. **(F)** Western blot of VSV-G-tagged EF_B0059 and Orf65/Orf65^L32F^ in *E. coli* strains from (E) 30 minutes post-induction. A non-specific band from the PAGE of these samples stained with Coomassie Brilliant Blue is used as a loading control (load). Data are representative of three independent experiments. **(G)** Western blot analysis of the Co-immunoprecipitation of VSV-G-tagged Orf65 or Orf65^L32F^ with FLAG-tagged EF_B0059. Tagged proteins were co-expressed in *E. coli* for 30 minutes upon which cells were lysed and FLAG-EF_B0059 was precipitated using anti-FLAG antibody conjugated to agarose beads. Data are representative of three independent experiments. Source data are provided as a Source Data file.

To understand how this phage protein prevents restriction by EF_B0059 we next used Alphafold2 to generate a high confidence model of Orf65^L32F^ (Fig. 3C and S5B)^58–60^. Orf65 is predicted to be a small protein consisting of two globular domains connected by a short linker. Structural alignments to proteins of known structure identified minimal overall similarity to characterized proteins, although the N-terminal domain within which the L32F mutation is localized bears some structural resemblance to SH3 domains of several eukaryotic proteins with the top scoring hit being the SH3 domain of myosin IB from *Entamoeba histolytica* (rmsd 2.3 Å, 9% sequence identity). In bacteria, SH3-like domains bind diverse ligands and molecules in a variety of unrelated proteins and thus this similarity provides minimal functional insight into how Orf65 confers resistance to EF_B0059^61^. However, in examining the predicted structure of Orf65, we also noted the presence of an unusually high number of acidic residues (21.3%) that are found on the surface of the protein resulting in a large electronegative surface potential (Fig. 3D). This high degree of surface electronegativity is reminiscent of the anti-restriction protein Ocr of phage T7. Ocr is a negatively charged protein (29% acidic amino acids) whose structure mimics B-DNA, which facilitates its binding and inactivation of Type I restriction enzymes by competing with their DNA substrates^62,63^. Although the predicted structure of Orf65^L32F^ does not resemble Ocr or mimic the structure of B-DNA, its similar electronegative surface properties led us to hypothesize that it may function as a direct inhibitor of EF_B0059.

If our hypothesis is correct, we would expect Orf65^L32F^ to inhibit the activity of EF_B0059 in the absence of any additional phi47 or *E. faecalis* encoded factors. To test this, we heterologously co-expressed EF_B0059 with either wildtype Orf65 or the Orf65^L32F^ variant in *E. coli* XL1-Blue. Consistent with our phi47 infection data in *E. faecalis*, expression of the Orf65^L32F^ variant but not the wildtype protein restored *E. coli* growth when co-expressed with EF_B0059 (Fig. 3E and 3F). Next, to determine if the observed Orf65^L32F^ inactivation is dependent on a physical interaction with EF_B0059, we immunoprecipitated FLAG-tagged EF_B0059 from cells co-expressing either VSV-G-tagged Orf65 or VSV-G-tagged Orf65^L32F^ (Fig. 3G). Western blotting of the precipitated fractions showed that wild-type EF_B0059 co-precipitates with a small amount of Orf65, which our co-expression data indicate is insufficient to alleviate the toxic effects of EF_B0059 on *E. coli* (Fig. 3E). However, we found that Orf65^L32F^ was substantially more enriched than the wildtype protein upon immunoprecipitation suggesting that the inhibitory properties of this variant are due to enhanced binding to EF_B0059. Importantly, in the absence of EF_B0059, neither Orf65 nor Orf65^L32F^ precipitate with α-FLAG conjugated agarose, confirming that the observed interactions are specific (Fig. S5C). Overall, these data suggest that while the wildtype Orf65 interacts with EF_B0059, the L32F mutation likely increases its binding affinity. Interestingly, mutation of leucine 32 to alanine or to other non-phenylalanine aromatic residues does not result in Orf65 variants capable of binding and inactivating EF_B0059, suggesting that the presence of phenylalanine at this position specifically stabilizes the Orf65^L32F^-EF_B0059 complex (Fig. S4E and S5D). Taken together, our data indicate that phi47 encodes a small, negatively charged protein that can evolve to inhibit the activity of EF_B0059. Furthermore, protein-protein interaction studies demonstrate that this inhibition is mediated by the direct binding of Orf65 to EF_B0059 and that this interaction is strongly influenced by a phenylalanine substitution at position 32 of Orf65.

### Orf65 represents a family of phage-encoded proteins that inhibit type IV restriction enzymes

The isolation of phage mutants that can infect phi47-resistant *E. faecalis* pTEF2 revealed that phi47 can evolve to overcome the restriction activity of EF_B0059. It achieves this through a single point mutation in the small protein Orf65, which promotes the binding and inactivation of EF_B0059. While wildtype Orf65 also interacts with EF_B0059, it does so with qualitatively lower affinity and does not display the same inhibitory effect as Orf65^L32F^. Intriguingly, sequence comparisons between Orf65 homologues found in the genomes of enterococcal phages shows that phenylalanine naturally occurs at position 32 in the *E. faecalis* phage SDS2, suggesting that phages related to phi47 may have previously evolved to overcome restriction by EF_B0059 and/or homologs of this TIV-RE (Fig. 4A). This observation suggests that the L32F substitution we isolated may be representative of an ongoing evolutionary arms race between phages that encode *orf65* and the hosts that use EF_B0059 and related proteins to restrict their growth. To probe this experimentally, we first asked if Orf65 from phage SDS2, which contains 84% sequence identity to Orf65 from phi47, can inactivate EF_B0059. However, owing to factors unrelated to the activity of EF_B0059, we found that phage SDS2 was unable to infect the phi47-susceptible strain V19 or the phi47-resistant strain V583. Further, we found that over-expression of the SDS2 Orf65 homolog is toxic to *E. coli* thereby precluding us from testing if it can inactivate EF_B0059 in this heterologous expression system (Fig. S6A). However, if EF_B0059 and Orf65 are co-evolving together as an effector-inhibitor pair, we reasoned that the wildtype Orf65 protein is likely already effective at inhibiting an EF_B0059-related restriction enzyme without requiring any amino acid substitutions. To test this hypothesis, we searched a database of human intestinal bacterial isolate genomes using the EF_B0059 sequence to identify candidate homologues^64^. As expected, genes from enterococci and related bacteria were enriched in this context. From this search, we identified two predicted TIV-REs including GC1503 from *Clostridium sporogenes,* and GC874 from *Leuconostoc mesenteroides*. These homologues share 57% sequence similarity to one another and approximately 40% sequence similarity to EF_B0059. Despite these relatively low sequence similarities, comparison of the Alphafold2 predicted structures of these proteins with that of EF_B0059 suggests that all three enzymes are structurally very similar (Fig. S7).

**Figure 4.**
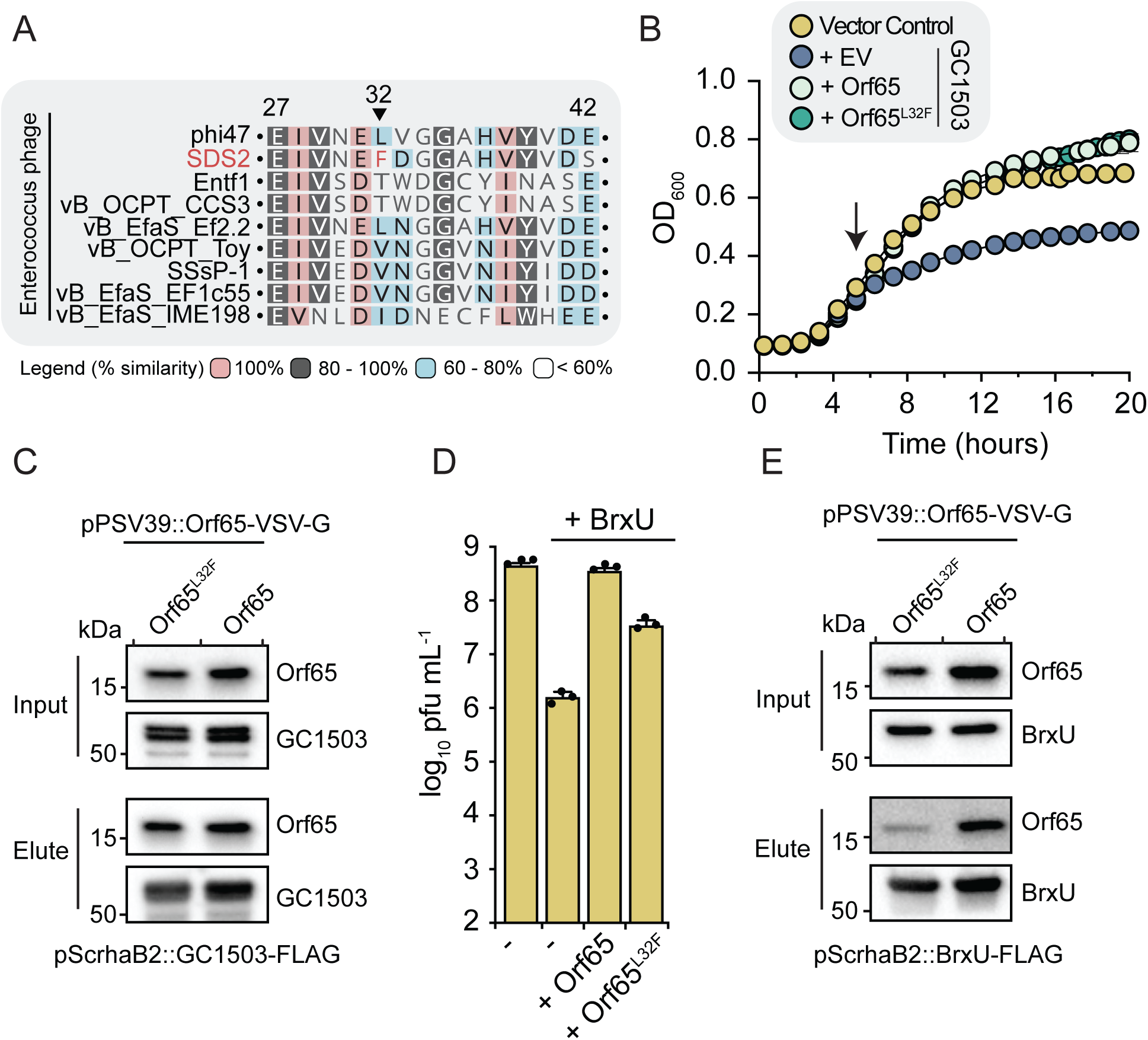
Wild-type Orf65 inactivates distantly related type IV restriction enzymes. **(A)** Multiple sequence alignment of residues 25-42 from Orf65 homologs identified in the genomes of 9 closely related enterococcal phages. The residues are colored according to their percent similarity at that site. **(B)** Growth curves of *E. coli* expressing GC1503 and either Orf65 or Orf65^L32F^, or empty vector. Arrow indicates when protein expression was induced. Data represent mean ± s.d. of three technical replicates. **(C)** Western blot analysis of the Co-immunoprecipitation of VSV-G-tagged Orf65 or Orf65^L32F^ with FLAG-tagged GC1503. Tagged proteins were co-expressed in *E. coli* for 30 minutes upon which cells were lysed and FLAG-tagged EF_B0059 was precipitated using anti-FLAG antibody conjugated to agarose beads. Data are representative of three independent experiments. **(D)** Outcome of coliphage infection measured by PFU/mL in *E. coli* strain XL-1 blue co-expressing BrxU and either Orf65 or Orf65^L32F^. Data represent mean ± s.d. of three technical replicates. **(E)** Outcome of Co-immunoprecipitation as in (C) but with FLAG-tagged BrxU. Source data are provided as a Source Data file.

Heterologous expression of these proteins in *E. coli* demonstrated that like EF_B0059, GC1503 is toxic to *E. coli*, whereas GC874 is not, suggesting GC1503 functions more similarly to EF_B0059 (Fig. S6B). Using this same assay, we asked if the co-expression of GC1503 with either Orf65 or Orf65^L32F^ restores *E. coli* growth. Remarkably, and in contrast to EF_B0059, we found that both wildtype and the L32F mutant of Orf65 were able to restore *E. coli* viability (Fig. 4B and S6C). Moreover, co-immunoprecipitation experiments using FLAG-tagged GC1503 revealed that wildtype Orf65 forms a complex with GC1503 that is comparable in abundance to that of the L32F mutant containing complex (Fig. 4C). These data indicate that although position 32 of Orf65 from phi47 is essential to stabilize its interaction with EF_B0059, it has no effect on its ability to inhibit GC1503. Indeed, each of the L32 variants (Trp, Tyr, and Ala) that we originally tested with EF_B0059 and were found to be non-protective are still protective against GC1503 activity (Fig. S6D). Therefore, the molecular mechanism of Orf65 binding to EF_B0059 homologous proteins appears to arise from multiple protein-protein contacts whose significance at the individual amino acid level differs depending on the TIV-RE in question.

We next asked if the inhibitory activity of Orf65 is generalizable to more divergent TIV-REs. To this end, we tested if the expression of Orf65 or its L32F variant interferes with the activity of BrxU, a recently characterized TIV-RE from the Gram-negative bacterium *Escherichia fergusonii,* which shares only 26% sequence similarity with EF_B0059^55^. Remarkably, expression of either Orf65 or Orf65^L32F^ interfered with the ability of BrxU to restrict the infection of *E. coli* by a waste-water isolated coliphage with wildtype Orf65 exhibiting a greater inhibitory effect than Orf65^L32F^ (Fig. 4D). In accordance with these data, co-immunoprecipitation experiments with FLAG-tagged BrxU found that both wildtype and Orf65^L32F^ interact with BrxU. However, in contrast to EF_B0059, the L32F mutation had a negative effect on the capacity of Orf65 to bind BrxU (Fig. 4E).

Together, our data support the conclusion that Orf65 of phi47 encodes an anti-TIV-RE factor. The results of our *E*. *coli* toxicity data in combination with our phage evolution experiments suggest that these phage proteins retain the ability to bind and inactivate diverse TIV-REs, but can evolve to become efficient inhibitors upon selection by a particular TIV-RE. Due to this defined role in overcoming anti-phage defense, we propose to name Orf65 type IV restriction inhibiting factor A (TifA).

## DISCUSSION

The need for alternative therapies to combat MDR bacterial infections has renewed interest in understanding how bacteria defend against phage infection. The majority of this research has prioritized the discovery and characterization of antiphage defense systems in model bacteria such as *E. coli* and *Bacillus subtilis,* with minimal focus on systems found in medically relevant and less-genetically tractable bacteria^20,21,25,65,66^. Moving forward there is a need to prioritize the characterization of such systems in bacteria that are responsible for MDR infections that will ultimately be the targets of phage therapy. In this study, we identify a previously unknown TIV-RE encoded on the enterococcal plasmid pTEF2 that limits phi47 phage replication. phi47 can overcome this restriction via a single point mutation of leucine to phenylalanine (L32F) in the previously uncharacterized protein Orf65, which herein has been renamed TifA. This mutation results in a gain-of-function inhibitory protein-protein interaction between TifA and EF_B0059. Intriguingly, EF_B0059 inhibition strictly depends on a phenylalanine at position 32 and is lost when other aromatic amino acids are substituted in this position. However, wildtype TifA binds and neutralizes diverse TIV-REs, suggesting that TifA represents a broadly active inhibitor of TIV-REs that undergoes adaptive evolution.

pTEF2 belongs to a family of plasmids known as pheromone responsive plasmids (PRPs) that can be mobilized among enterococcal species. PRPs frequently carry genes required for antimicrobial resistance and pathogenicity and thus play a crucial role in virulence^67^. Here, we show these plasmids also contribute to the ongoing battle between enterococci and their phage parasites by encoding dedicated antiphage defense systems. Considering the pervasiveness of phage predation within complex microbial ecosystems such as the mammalian gut combined with the proficient PRP-mediated genetic transfer observed in these contexts, we speculate that PRPs contribute significantly to the ability of the enterococci to counter phage threats, potentially serving both as a reservoir for and mechanism to mobilize antiphage defense systems^68,69^.

We further discovered that within pTEF2, EF_B0059 is found in an operon with EF_B0058, which is predicted to encode a serine recombinase. Such recombinases have well-established roles as mediators of genotypic diversity and population-level fitness in bacteria^70^. Recent studies indicate that recombinases are frequently found in phage defense islands, and although their direct contribution to the mobilization of antiphage defenses remains unconfirmed, they are presumed to aid in the diversification of defense islands via homologous recombination^14^. Indeed, so-called defense islands are commonly used to discover novel antiphage systems, however, the mechanisms that drive their genetic variation and mobility are poorly understood. It will be of interest in future studies to probe if and how such recombinases contribute to the diversification of antiphage defense systems in the enterococci and more broadly across bacterial taxa.

We found that EF_B0059 provides *E. faecalis* resistance to infection by the lytic phage phi47. The precise mechanism by which EF_B0059 prevents phi47 replication is unknown, however, we found that expression of EF_B0059 is toxic to *E. coli* and that this toxicity is abrogated upon mutation of conserved active site residues in either the N-terminal DUF262 or C-terminal DUF1524 domains. These data suggest that EF_B0059 functions similarly to other enzymes that harbor these domains such as the TIV-REs GmrSD and BrxU, which target and degrade DNA methylated at cytosine bases. Interestingly, however, we also found that EF_B0059 remains toxic to an *E. coli* strain that is unable to methylate its DNA. In line with this finding, a sequence analysis of the phi47 genome failed to identify genes with homology to known DNA modifying enzymes, suggesting phi47 does not modify its DNA. This notion is further reinforced by the mutations identified in our isolation of escape phages that evade EF_B0059 restriction. If phi47 encoded DNA modifying enzymes that sensitize it to restriction by EF_B0059, we would expect to isolate phage mutants for any such genes. The lack of identification of such genes in the phi47 genome in addition to the LC-MS analysis of phi47 DNA showing no obvious modifications, suggests that phi47 does not directly modify its DNA. Together, these observations indicate that EF_B0059 targets unmodified DNA, distinguishing it from previously characterized TIV-REs. This raises the obvious question of how EF_B0059 differentiates between host and phage DNA. One possible explanation is that EF_B0059 functions more akin to classical restriction-modification systems in which functionally and genetically paired modification systems modify the host genome at specific sites that block cleavage by their cognate restriction endonuclease. This explanation seems unlikely given that the 14 genes we identified as being unique to pTEF2 compared to EF_B0059-deficent pCF10, are not predicted to encode proteins with the capacity to modify DNA. A more plausible explanation is that EF_B0059 recognizes a feature of the phage genome that is unrelated to a specific base modification, such as a unique sequence motif or higher-order DNA structure. For example, It has been previously reported that Wadjet systems cleave closed-circular DNA^71^. Moreover, EF_B0059 activity could be stimulated by an unknown signal that is present during phage infection, perhaps related to the entry of the phage genome into the cell or the early stages of its replication. Future studies examining how phages lacking a TifA homolog are able to escape EF_B0059-dependent restriction may answer some of these questions and shed light on how this defense system discriminates between self and non-self DNA.

It is important to note that protein-based antirestriction mechanisms, including those that target TIV-REs have been described previously. Indeed, the discovery of GmrSD in *E. coli* as a modification-dependent restriction enzyme that prevents T4 phage replication also coincided with the characterization of the T4 phage internal <u>p</u>rotein I (IPI) as an inhibitor of GmrSD. IPI is packaged into the T4 capsid and upon infection is released into the host cell where it protects the phage DNA from degradation by binding to and inhibiting the enzymatic activity of GmsrSD^53,72^. Our data indicate that like IPI, TifA inhibits EF_B0059 through a direct binding mechanism. However, IPI and TifA share neither sequence nor structural homology indicating that these TIV-RE inhibitors evolved independently. Intriguingly, the independent emergence of counter-defense proteins like IPI and TifA, which target the same family of defense systems, might provide insight into why phage genomes are replete with hypothetical genes that lack homology to other genes beyond those found in closely related phages. It is possible that many of these encode species-specific counter defense proteins that have independently evolved to counteract related bacterial defense mechanisms. Our discovery of TifA adds to a growing list of recent phage-bacteria coevolution studies showing that many of these small cryptic phage proteins function specifically to counter antiphage defense^73,74^.

Our data show that the wild-type TifA can bind and disrupt the activity of GC1503 and BrxU, whereas a single missense mutation provides it with the capacity to inhibit EF_B0059. Given their divergent sequences, these findings raise the question as to how TifA is able to interact with and inhibit such diverse TIV-REs. We suspect that an explanation may lie in unique biochemical characteristics of TifA. Specifically, our structural modeling indicates that TifA has a highly electronegative surface that likely facilitates its binding to the electropositive regions of TIV-REs that are involved in DNA binding. Although the details underpinning DNA recognition in TIV-REs are incomplete, recent biochemical and structural studies of BrxU demonstrated that its DNase activity is linked to its homodimerization, which results in the formation of a large electropositive cleft that the authors posit is well suited for DNA binding. In the case of TifA, adaptive mutations that enhance its binding to a particular TIV-RE would be selected for as improved inhibitory activity would result in successful phage infections. Such a mechanism of inhibition would explain why we observed a weak interaction between EF_B0059 and wild-type phi47 TifA that is enhanced by the L32F mutation identified in the phi47 escape mutants.

Competitive inhibition models have been established for several antirestriction proteins targeting other types of restriction enzymes. For example, the T7 phage protein Ocr competitively inhibits multiple diverse Type I restriction enzymes by mimicking the structure and chemical properties of B-form DNA^63^. As we suspect is the case for TifA, selection for point mutations in Ocr results in a more efficient inhibitor of specific type I restriction enzymes^62^. Although our data suggest that Ocr and TifA may be functionally analogous in their capacity to inhibit multiple restriction enzymes, the predicted structure of TifA does not appear to explicitly mimic the charge distribution of DNA in any obvious way and thus we refrain from classifying TifA as a bona fide DNA mimic. Indeed, further biochemical and structural investigations are required to understand how TifA inhibits TIV-REs. Nevertheless, our data suggest that like DNA-mimic antirestriction proteins, TifA likely provides phi47 and related phages with a robust antiphage defense strategy, allowing these phages to adapt to and inhibit diverse TIV-REs.

## METHODS

### Bacterial strains, bacteriophages, and growth conditions

All bacterial and bacteriophage strains used in this study can be found in Table S2. *E. faecalis* strains were grown with aeration in Brain Heart Infusion (BHI, Becton Dickinson) broth or on BHI agar at 37°C. To maintain pLZ12A, pMSP3545, pGCP123 plasmids cultures were supplemented with 15 µg/mL chloramphenicol, 20 µg/ml erythromycin, and 500 µg/mL kanamycin, respectively. For induction of the CRISPRi plasmids, cells were grown with 50 ng/mL nisin (MilliporeSigma). Phage plaque assays were performed using Todd-Hewitt broth (THB, Becton Dickinson) agar supplemented with 10 mM MgSO_4_ as described elsewhere^29^. To generate strains of *E. faecalis* carrying pTEF2 and its derivatives, we performed pheromone responsive plasmid matings. V19 pTEF2 and derivatives were selected with 7.5 µg/mL vancomycin and 200 µg/mL spectinomycin. JH2-2 pTEF2 and derivatives were selected with 50 µg/mL rifampicin and 200 µg/mL spectinomycin. The construction of EF_B0058 and EF_B0059 mutant strains of *E. faecalis* followed a previously published protocol for allelic exchange^75^. Primers used to generate deletion constructs that were cloned into pLT06 can be found in Table S3. *Escherichia coli* and *Escherichia fergusonii* were grown in Lennox lysogeny broth (LB, Fisher Scientific) with aeration or on LB agar at 37°C. When appropriate, cultures were supplemented with 15 µg/mL chloramphenicol, 20 µg/ml erythromycin, 50 µg/mL kanamycin, 200 μg/mL trimethoprim, 15 μg/mL gentamicin, 0.5 mM IPTG, and 0.2% w/v L-rhamnose. *Clostridium sporogenes* and *Leuconostoc mesenteroides* were grown on BHI broth and agar supplemented with 0.5 g/L L-cysteine, 10 mg/L hemin, and 1 mg/L vitamin K at 37°C in an anerobic chamber (5% CO_2_, 5% H_2_, 90% N_2_; Shel Labs).

### Plasmid construction

A detailed list of the primers and plasmids used in this study can be found in Table S2. For DNA amplification by PCR, *E. faecalis, E fergusonii, L. mesenteroides, C. sporogenes,* and phi47 genomic DNA was purified using a ZymoBIOMICS DNA Miniprep Kit (Zymo Research) or a previously published protocol for phenol/chloroform extraction^29^. PCR reactions used to create DNA fragments for allelic exchange were performed using KOD Hot Start DNA Polymerase (EMD Millipore) and cloned into pLT06^75^. phi47 *orf65* (*tifA*) and *orf65*-L32F were PCR amplified using Q5 master mix (New England Biolabs) and cloned into pLZ12A, a derivative of the shuttle vector pLZ12 carrying the *bacA* promoter (41). CRISPRi sgRNA pGCP123 plasmids were designed and cloned following previously published protocols^50^. Plasmids were transformed into *E. faecalis* by electroporation. Briefly, *E. faecalis* was grown in 100 ml of BHI broth to an OD_600_ of 0.3. The cells were washed once with 10 ml of ice-cold 10% glycerol and resuspended in 500 µl of lysozyme solution (30 μg/ml lysozyme, 20% sucrose, 10 mM EDTA, 50 mM NaCl and 10 mM Tris-HCl, pH 8.0) and incubated at 37°C for 20 min. The cells were washed four times with 500 µl of ice-cold electroporation buffer (500 mM sucrose and 10% glycerol). Cell pellets were resuspended in 250 µl of ice-cold electroporation buffer and 0.5 – 1.0 µg of plasmid DNA was electroporated into 50 µl of the *E. faecalis* suspension using a 0.2 mm cuvette at 2.5k V, 200 Ω, and 25 µF. Plasmids used in *E. coli* include pSCrhaB2-CV, a low-copy (20-30 copies per cell), tightly regulated expression vector containing a rhamnose-inducible promotor used for the expression of EF_B0059 and other TIV-REs and pPSV39-CV, a high-copy expression vector containing a *lac*UV5 promoter used for the expression of TifA^76–78^. All expression plasmids were constructed using standard restriction enzyme-based cloning procedures. Reagents used for cloning, including restriction enzymes, T4 DNA ligase, and Phusion DNA polymerase were obtained from New England Biolabs. Plasmids were transformed into chemically competent *E. coli* by heat-shock and transformants were subsequently isolated by selection with the appropriate antibiotics.

### Genomic comparisons of *E. faecalis* strains

The chromosomes of *E. faecalis* SF28073 (NZ_CP060804.1) and V583 (NC_004668.1) were obtained from NCBI and aligned using the MAUVE multiple genome alignment tool^79^. To compare plasmid sequences between *E. faecalis* strains, sequences of the plasmids pTEF1 (NC_004669.1), pTEF2 (NC_004670.1), pTEF3 (NC_004671.1), pSF1 (NZ_CP060801.1), pSF2 (NZ_CP060802.1), and pSF3 (NZ_CP060803.1) were obtained from NCBI and aligned using the clinker alignment tool^80^.

### Pheromone responsive plasmid mating

For pheromone responsive plasmid conjugation, 150 µL of overnight *E. faecalis* cultures were subcultured into 5 mL of BHI broth and grown for 90 minutes with aeration at 37°C. Donor and recipient bacteria were mixed at a 1:1 ratio in a volume of 1 mL and applied to a 0.45-micron filter. The filter was removed from the housing and placed on a BHI agar plate with the bacteria side of the filter facing up. These plates were incubated overnight at 37°C. The following day, the filter was removed from the plate and placed in 5 mL of sterile PBS and vortexed to dislodge the bacteria from the filter. Serially diluted bacteria were plated on BHI agar containing antibiotics as described above to select for recipient cells carrying the plasmid and were grown at 37°C for no more than 48 hrs. Potential transconjugants were screened via colony PCR to confirm the presence of the plasmid. To generate *E. faecalis* V19 pTEF2 and JH2-2 pTEF2 the donor was *E. faecalis* OG1RF pTEF2-specR^40^. To generate *E. faecalis* V19 pTEF1, *E. faecalis* OG1SSp pTEF1 was used as the donor^40^.

### Growth curves

For broth culture infections of *E. faecalis* with phi47, 15 mL of THB broth was inoculated with *E. faecalis* and grown to an OD_600_ of 0.3. Cultures were infected with phi47 at a multiplicity of infection of either 1 or 10, as indicted, and supplemented with 10 mM MgSO_4_. Cultures were incubated at room temperature for 5 minutes, then moved to 37°C with aeration. Phage production was quantified by removing 100 µL of the culture, adding 33.3 µL of chloroform, and quantifying the number of phages using plaque assays^38^. For the *E. coli* growth curves, overnight cultures containing the indicated plasmids were diluted 1:100 into LB broth supplemented with 200 μg/mL trimethoprim and 15 μg/mL gentamicin. For each culture, 200 μL was seeded into a 96-well plate in triplicate. Growth was measured at 15-minute intervals until an OD_600_ of 0.4 was reached, upon which protein expression was induced by the addition of either 0.2% w/v L-rhamnose or 0.2% w/v L-rhamnose and 0.5 mM IPTG. For the infection of *E. coli* cells expressing BrxU and Orf65 with coliphage, overnight cultures were diluted 1:100 in LB and grown to an OD_600_ of 0.5. Expression of Orf65 and/or BrxU was induced by the addition 0.5 mM IPTG and 0.2% w/v L-rhamnose, respectively. After 30 minutes, 500 μL of *E. coli* was mixed with 5 mL of semi-solid LB (0.75% w/v agar) supplemented with 0.5 mM IPTG and/or 0.2% L-rhamnose and poured onto an 1.5% w/v LB agar plate. The agar overlay was allowed to dry and 5 μL of serially diluted coliphages were spotted on top. After overnight incubation at 37 °C, the plaques were quantified and compared between inducing conditions.

### Reverse transcription-PCR

cDNA was generated from 1 µg of RNA using qScript cDNA SuperMix (QuantaBio) and the cDNA was purified using a QIAquick PCR Purification Kit (Qiagen). 2 µL of cDNA was used in a PCR reaction with primers specific to EF_B0058 and EF_B0059 (Table S3). The resulting PCR products were run on a 1% w/v agarose gel to visualize PCR bands.

### Phage adsorption assay

*E. faecalis* cultures grown in BHI were pelleted at 3220 × *g* for 10 minutes and resuspended to 10^8^ CFU/mL in SM-plus buffer (100 mM NaCl, 50 mM Tris-HCl, 8LJmM MgSO_4_, 5LJmM CaCl_2_ [pH 7.4]). Phages were added at an MOI of 0.1 and incubated at room-temperature without agitation for 10 minutes. The bacteria-phage suspensions were centrifuged at 24,000 × *g* for 1 minute and the supernatant was collected to determine the number of unbound phages using a plaque assay. SM-plus buffer supplemented with phage only was used as a negative control.

### Western blotting

To analyze the expression of EF_B0059, its active site mutants, and Orf65, 5 mL cultures of *E. coli* containing the indicated plasmids were grown to an OD_600_ of 0.5, upon which protein expression was induced by the addition of 0.2% w/v L-rhamnose and 0.5 mM IPTG. The cells were pelleted by centrifugation at 7,800 × *g* after 30 minutes of protein expression. Cell pellets were resuspended in 50 μL of PBS and mixed 1:1 with Laemmli buffer and boiled at 95°C for 10 minutes. Samples were centrifuged at 16,000 × *g* and 5 μL was loaded onto a 10% SDS PAGE gel. The gel was run for 15 minutes at 95 V, followed by 40 minutes at 195 V, and total protein was wet transferred to a nitrocellulose membrane using a mini Trans-Blot electrophoretic system (Bio-Rad). After the transfer, the blot was blocked in TBS-T supplemented with 5% w/v non-fat milk, shaking gently at room temperature for 1 hour. The blot was washed three times with TBS-T and primary antibody - either anti-VSV-G (MilliporeSigma, V4888) or anti-FLAG (MilliporeSigma, F7425) - was added to the blot at a 1:5000 dilution followed by an overnight incubation shaking gently at 4°C. The blot was washed three times with TBS-T, followed by a 1-hour incubation with secondary antibody (anti-rabbit HRP conjugate). After three more washes with TBS-T, the blot was developed by the addition of Clarity Max ECL substrate (Bio-Rad) and chemiluminescence was visualized using a ChemiDoc Touch Imaging System (Bio-Rad). Images of uncropped and unprocessed gels are presented in the supplementary information file (Fig. S8).

### CRISPRi screening

Using previously described CRISPRi constructs for *E. faecalis*^50^, each gene unique to pTEF2 was knocked down. In brief, pMSP3545-dCas9_Str_ was leveraged to express the dCas9 to inhibit gene expression, while pGCP123 carried each of the guide RNAs (sgRNAs) to facilitate targeted inhibition. sgRNA oligonucleotides were designed against pTEF2 open reading frames and targeted the non-template strand. sgRNA oligonucleotides were ordered from Twist Bioscience. All sgRNA oligonucleotides used can be found in Table S3. *E. faecalis* V19 pTEF2 with CRISPRi and sgRNA plasmids were cultured overnight in BHI broth containing 200 μg/mL kanamycin, 50 μg/mL erythromycin, and 50 ng/mL nisin. Overnight cultures were diluted to an OD_600_ of 0.1 in SM-plus buffer. 6 µL of 10-fold serial dilutions were spotted on THB agar plates supplemented with 10mM MgSO_4_, 200 μg/mL kanamycin, 50 μg/mL erythromycin, and 50 ng/mL nisin, with and without 1×10^7^ PFU/mL of phi47 embedded in the agar. Plates were incubated at 37°C for 48 hours and bacteria were enumerated. Relative viability was calculated by dividing the CFU of cells grown in the presence of phi47 by the CFU grown in the absence of phi47.

### LC-MS analysis of phi47 DNA

DNA isolated from phage phi47 was digested into nucleosides before being subjected to liquid chromatography-mass spectrometry (LC-MS) analysis. Approximately 1-2 µg phi47 DNA was treated with Nucleoside Digestion Mix (New England Biolabs) following the manufacturer’s protocol at 37°C for >2h. The resulting mixture of nucleosides were filtered through a hydrophilic PTFE 0.2 µm centrifugal filter, and the filtrate was subjected to the reverse phase LC-MS for nucleoside separation and MS detection. LC-MS was performed on an Agilent 1290 Infinity II UHPLC-MS system equipped with a G7117 Diode Array Detector and a LC/MSD XT G6135 Single Quadrupole Mass Detector. Chromatography performed with the instrument was on a Waters XSelect HSS T3 C18 column (4.6 × 150 mm, 3 µm particle size) operated at a flow rate of 0.5 mL/min with a binary gradient mobile phase consisting of 10 mM ammonium acetate (pH 4.5) and methanol. The course of chromatography was monitored at 260 nm. MS was operated in both positive (+ESI) and negative (-ESI) electrospray ionization modes. MS was performed with a capillary voltage of 2500 V at both modes, a fragmentation voltage of 70 V, and a mass range of m/z 100 to 1000. Agilent ChemStation software was used for the primary LC-MS data processing.

### Structural modeling and comparisons

Protein structures were modelled using AlphaFold2 within Colabfold (V.1.5.2) that is part of Google Collaboratory. Predictions were run using the default parameters^60^. The resulting structures were visualized, and electrostatic surface potential was mapped using ChimeraX^81^. Structural comparisons and RMSD calculations were performed using the DALI webserver^82^.

### Identification of bacteria encoding GC1503 and GC874

To identify homologs of EF_B0059, we searched a local database containing the Surette lab whole genome collection of diverse human bacterial phyla^64^. For this, the amino acid sequence of EF_B0059 was searched against this database using tblastn in BLAST v2.13.0+, with default settings and the ‘-t 6’ option for generating tabular results^83^. The taxonomy of the genomes was identified using GTDB-Tk^2^ v202.0, and the results were merged with the BLAST hits in R version 4.0.3^84^.

### Isolation of *E. coli* bacteriophages from wastewater

Bacteriophages were isolated from wastewater as previously described with minor alterations^55^. In brief, 5 mL of wastewater was centrifuged at 9,000 × *g* and then filtered through a 0.22 μM filter. The cleared wastewater was mixed with 5 mL of LB and inoculated with 500 μL of an overnight *E. coli* culture. After a 24-hour incubation shaking at 37°C, 2 mL aliquots were removed from cultures that showed evidence of phage propagation (complete cell lysis), and centrifuged at 12,000 × *g*. The supernatant was filtered through a 0.22 μM filter to remove any remaining bacterial cells. The filtered supernatant was serially diluted in SM-plus buffer and 10 µL was added to 500 µL of an overnight *E. coli* culture. This phage-bacterial mixture was then added to 5 mL of semi-solid LB (0.75% w/v agar), mixed gently, and overlayed onto an LB agar plate. After overnight incubation, the resulting plaques were isolated. Agar plugs of individual plaques were used to inoculate cultures of *E. coli* grown to mid-log phase as determined by OD_600_ measurements. Upon lysis of the *E. coli* cultures, the resulting phages were purified as described above. The above infection protocol was repeated until a uniform plaque morphology was obtained. Using this approach, we isolated 2 phages from 8 unique wastewater samples for a total of 16 isolated coliphages. Subsequent plaque assays found that the replication of six of these phages were restricted by BrxU to varying degrees. From these six, we used the phage that displayed the largest degree of replication inhibition, which we term JCW01, to assay if co-expression of Orf65/Orf65^L32F^ interfered with BrxU-dependent restriction.

### Sequencing and genomic analysis of pTEF2 escape phi47 isolates

phi47 genomic DNA was sequenced using the Illumina NextSeq 2000 platform to 300 Mbp depth at SeqCenter (Pittsburgh, PA). The resulting reads were mapped to the phi47 genome (NCBI accession ON086985.1) and SNPs were identified using CLC Genomics Workbench v20 (Qiagen) with a minimum coverage of 15, a minimum frequency of 30%, and ploidy of 0.

### Co-immunoprecipitation assays

To probe the interaction between TifA and a given TIV-RE, 50 mL cultures of *E. coli* were grown to an OD_600_ of 0.5 upon which expression of FLAG-tagged TIV-RE and VSV-G-tagged Orf65 were induced with the addition 0.2% w/v L-rhamnose and 0.5 mM IPTG, respectively. After 30 minutes the cells were pelleted and resuspended in 2 mL of lysis buffer (150 mM NaCl, 20 mM Tris-HCl pH 7.5) and lysed by sonication. The cell lysates were cleared by centrifugation at 16,000 × *g* for 30 minutes at 4 °C. Cleared cell lysates were incubated with 50 μL of anti-FLAG agarose beads (Genscript, L00425) for 2 hours at 4 °C using a tube rotator. The beads were washed three times with 10 mL of lysis buffer and bound protein was eluted by boiling in 50 μL of Laemmli buffer. 5 μL of the elution was subjected to SDS-PAGE and western blotting as described above using anti-FLAG and anti-VSV-G as the primary antibodies as described above.

## DATA AVAILABILITY

The DNA sequencing reads generated in this study can be found at the European Nucleotide Archive under project accession PRJEB68205 [https://www.ebi.ac.uk/ena/browser/view/PRJEB68205]. Raw data for plotted figures are provided in the Source Data file. Raw images for immunoblots are provided in the Supplementary Information file.

## Supporting information

Combined Supplementary Figures and Tables

## ACKNOWLEDGEMENTS

We would like to thank Dr. Michael Gilmore for providing *E. faecalis* OG1RF pTEF2-specR and *E. faecalis* OG1SSp pTEF1, Dr. Katrine Whiteson for sharing phage SDS2, Dr. Kimberly Kline for providing CRISPRi genetic tools, Dr. James Samuelson and Dr. Elisabeth Raleigh for supplying the *E. coli* strain ER2796, and Dr. Michael Surette for sharing *Clostridium sporogenes* and *Leuconostoc mesenteroides* strains. We also thank Dr. Lindsey Marmont for her constructive feedback on the manuscript.

## AUTHOR CONTRIBUTIONS

B.D., J.W., C. J., N.B., and S. A. conceptualized the project. C.J. and N.B. performed the majority of experiments. S. A., G. A., S. M., and Y-J. L. assisted with experiments. P. W. provided valuable insight for data analysis. C. J. and N. B. wrote the manuscript. B. D., J. W., and S. A. edited the manuscript. B. D., J. W., C. J., S. A., and G. A. acquired funding that supported the study.

## REFERENCES

1. Donskey CJ, Chowdhry TK, Hecker MT, Hoyen CK, Hanrahan JA, Hujer AM, Hutton-Thomas RA, Whalen CC, Bonomo RA, Rice LB. Effect of antibiotic therapy on the density of vancomycin-resistant enterococci in the stool of colonized patients. N Engl J Med. 343, 1925–1932 (2000). doi:10.1056/NEJM200012283432604

2. Ubeda C, Bucci V, Caballero S, Djukovic A, Toussaint NC, Equinda M, Lipuma L, Ling L, Gobourne A, No D, Taur Y, Jenq RR, van den Brink MRM, Xavier JB, Pamer EG. Intestinal microbiota containing *Barnesiella* species cures vancomycin-resistant *Enterococcus faecium* colonization. Infect Immun. 81, 965–973 (2013). doi:10.1128/IAI.01197-12

3. Suh GA, Lodise TP, Tamma PD, Knisely JM, Alexander J, Aslam S, Barton KD, Bizzell E, Totten KMC, Campbell JL, Chan BK, Cunningham SA, Goodman KE, Greenwood-Quaintance KE, Harris AD, Hesse S, Maresso A, Nussenblatt V, Pride D, Rybak MJ, Sund Z, van Duin D, Van Tyne D, Patel R. Considerations for the use of phage therapy in clinical practice. Antimicrob Agents Chemother. 66, e0207121 (2022). doi:10.1128/aac.02071-21

4. Strathdee SA, Hatfull GF, Mutalik VK, Schooley RT. Phage therapy: From biological mechanisms to future directions. Cell. 186, 17–31 (2023). doi:10.1016/j.cell.2022.11.017

5. Kortright KE, Chan BK, Koff JL, Turner PE. Phage therapy: A renewed approach to combat antibiotic-resistant bacteria. Cell Host Microbe. 25, 219–232 (2019). doi:10.1016/j.chom.2019.01.014

6. Sulakvelidze A, Alavidze Z, Morris JG. Bacteriophage therapy. Antimicrob Agents Chemother. 45, 649–659 (2001). doi:10.1128/AAC.45.3.649-659.2001

7. Langdon A, Crook N, Dantas G. The effects of antibiotics on the microbiome throughout development and alternative approaches for therapeutic modulation. Genome Med. 8, 39 (2016). doi:10.1186/s13073-016-0294-z

8. Moelling K, Broecker F, Willy C. A wake-up call: We need phage therapy now. Viruses. 10, 688 (2018). doi:10.3390/v10120688

9. Brives C, Pourraz J. Phage therapy as a potential solution in the fight against AMR: obstacles and possible futures. Palgrave Commun. 6, 1–11 (2020). doi:10.1057/s41599-020-0478-4

10. Cook R, Brown N, Redgwell T, Rihtman B, Barnes M, Clokie M, Stekel DJ, Hobman J, Jones MA, Millard A. INfrastructure for a PHAge REference Database: Identification of large-scale biases in the current collection of cultured phage genomes. Phage (New Rochelle). 2, 214–223 (2021). doi:10.1089/phage.2021.0007

11. Ofir G, Sorek R. Contemporary phage biology: From classic models to new insights. Cell. 172, 1260–1270 (2018). doi:10.1016/j.cell.2017.10.045

12. Hampton HG, Watson BNJ, Fineran PC. The arms race between bacteria and their phage foes. Nature. 577, 327–336 (2020). doi:10.1038/s41586-019-1894-8

13. Duncan-Lowey B, Kranzusch PJ. CBASS phage defense and evolution of antiviral nucleotide signaling. Curr Opin Immunol. 74, 156–163 (2022). doi:10.1016/j.coi.2022.01.002

14. Hochhauser D, Millman A, Sorek R. The defense island repertoire of the *Escherichia coli* pan-genome. PLoS Genet. 19, e1010694 (2023). doi:10.1371/journal.pgen.1010694

15. Makarova KS, Wolf YI, Snir S, Koonin EV. Defense islands in bacterial and archaeal genomes and prediction of novel defense systems. J Bacteriol. 193, 6039–6056 (2011). doi:10.1128/jb.05535-11

16. Doron S, Melamed S, Ofir G, Leavitt A, Lopatina A, Keren M, Amitai G, Sorek R. Systematic discovery of antiphage defense systems in the microbial pangenome. Science. 359, (2018). doi:10.1126/science.aar4120

17. Brouns SJ, Jore MM, Lundgren M, Westra ER, Slijkhuis RJ, Snijders AP, Dickman MJ, Makarova KS, Koonin EV, van der Oost J. Small CRISPR RNAs guide antiviral defense in prokaryotes. Science. 321, 960–964 (2008). doi:10.1126/science.1159689

18. Mojica FJ, Diez-Villasenor C, Garcia-Martinez J, Soria E. Intervening sequences of regularly spaced prokaryotic repeats derive from foreign genetic elements. J Mol Evol. 60, 174–182 (2005). doi:10.1007/s00239-004-0046-3

19. Tock MR, Dryden DT. The biology of restriction and anti-restriction. Curr Opin Microbiol. 8, 466–472 (2005). doi:10.1016/j.mib.2005.06.003

20. Xiong X, Wu G, Wei Y, Liu L, Zhang Y, Su R, Jiang X, Li M, Gao H, Tian X, Zhang Y, Hu L, Chen S, Tang Y, Jiang S, Huang R, Li Z, Wang Y, Deng Z, Wang J, Dedon PC, Chen S, Wang L. SspABCD-SspE is a phosphorothioation-sensing bacterial defence system with broad anti-phage activities. Nat Microbiol. 5, 917–928 (2020). doi:10.1038/s41564-020-0700-6

21. Goldfarb T, Sberro H, Weinstock E, Cohen O, Doron S, Charpak-Amikam Y, Afik S, Ofir G, Sorek R. BREX is a novel phage resistance system widespread in microbial genomes. EMBO J. 34, 169–183 (2015). doi:10.15252/embj.201489455

22. Pecota DC, Wood TK. Exclusion of T4 phage by the *hok*/*sok* killer locus from plasmid R1. J Bacteriol. 178, 2044–2050 (1996). doi:10.1128/jb.178.7.2044-2050.1996

23. Fineran PC, Blower TR, Foulds IJ, Humphreys DP, Lilley KS, Salmond GP. The phage abortive infection system, ToxIN, functions as a protein-RNA toxin-antitoxin pair. Proc Natl Acad Sci U S A. 106, 894–899 (2009). doi:10.1073/pnas.0808832106

24. Dy RL, Przybilski R, Semeijn K, Salmond GP, Fineran PC. A widespread bacteriophage abortive infection system functions through a Type IV toxin-antitoxin mechanism. Nucleic Acids Res. 42, 4590–4605 (2014). doi:10.1093/nar/gkt1419

25. Cohen D, Melamed S, Millman A, Shulman G, Oppenheimer-Shaanan Y, Kacen A, Doron S, Amitai G, Sorek R. Cyclic GMP-AMP signalling protects bacteria against viral infection. Nature. 574, 691–695 (2019). doi:10.1038/s41586-019-1605-5

26. Millman A, Bernheim A, Stokar-Avihail A, Fedorenko T, Voichek M, Leavitt A, Oppenheimer-Shaanan Y, Sorek R. Bacterial retrons function in anti-phage defense. Cell. 183, 1551–1561 e1512 (2020). doi:10.1016/j.cell.2020.09.065

27. Kronheim S, Daniel-Ivad M, Duan Z, Hwang S, Wong AI, Mantel I, Nodwell JR, Maxwell KL. A chemical defence against phage infection. Nature. 564, 283–286 (2018). doi:10.1038/s41586-018-0767-x

28. Burmeister AR, Fortier A, Roush C, Lessing AJ, Bender RG, Barahman R, Grant R, Chan BK, Turner PE. Pleiotropy complicates a trade-off between phage resistance and antibiotic resistance. Proc Natl Acad Sci U S A. 117, 11207–11216 (2020). doi:10.1073/pnas.1919888117

29. Duerkop BA, Huo W, Bhardwaj P, Palmer KL, Hooper LV. Molecular basis for lytic bacteriophage resistance in enterococci. mBio. 7, e01304–01316 (2016). doi:10.1128/mBio.01304-16

30. Gordillo Altamirano F, Forsyth JH, Patwa R, Kostoulias X, Trim M, Subedi D, Archer SK, Morris FC, Oliveira C, Kielty L, Korneev D, O’Bryan MK, Lithgow TJ, Peleg AY, Barr JJ. Bacteriophage-resistant *Acinetobacter baumannii* are resensitized to antimicrobials. Nat Microbiol. 6, 157–161 (2021). doi:10.1038/s41564-020-00830-7

31. Eugster MR, Morax LS, Huls VJ, Huwiler SG, Leclercq A, Lecuit M, Loessner MJ. Bacteriophage predation promotes serovar diversification in *Listeria monocytogenes*. Mol Microbiol. 97, 33–46 (2015). doi:10.1111/mmi.13009

32. Canfield GS, Chatterjee A, Espinosa J, Mangalea MR, Sheriff EK, Keidan M, McBride SW, McCollister BD, Hang HC, Duerkop BA. Lytic bacteriophages facilitate antibiotic sensitization of *Enterococcus faecium*. Antimicrob Agents Chemother. 65, e00143–00121, AAC.00143-00121 (2021). doi:10.1128/AAC.00143-21

33. Paul BG, Burstein D, Castelle CJ, Handa S, Arambula D, Czornyj E, Thomas BC, Ghosh P, Miller JF, Banfield JF, Valentine DL. Retroelement-guided protein diversification abounds in vast lineages of bacteria and archaea. Nat Microbiol. 2, 17045 (2017). doi:10.1038/nmicrobiol.2017.45

34. Rusinov IS, Ershova AS, Karyagina AS, Spirin SA, Alexeevski AV. Avoidance of recognition sites of restriction-modification systems is a widespread but not universal anti-restriction strategy of prokaryotic viruses. BMC Genomics. 19, 885 (2018). doi:10.1186/s12864-018-5324-3

35. Golovenko D, Manakova E, Tamulaitiene G, Grazulis S, Siksnys V. Structural mechanisms for the 5’-CCWGG sequence recognition by the N- and C-terminal domains of EcoRII. Nucleic Acids Res. 37, 6613–6624 (2009). doi:10.1093/nar/gkp699

36. Blower TR, Evans TJ, Przybilski R, Fineran PC, Salmond GP. Viral evasion of a bacterial suicide system by RNA-based molecular mimicry enables infectious altruism. PLoS Genet. 8, e1003023 (2012). doi:10.1371/journal.pgen.1003023

37. Samson JE, Belanger M, Moineau S. Effect of the abortive infection mechanism and type III toxin/antitoxin system AbiQ on the lytic cycle of *Lactococcus lactis* phages. J Bacteriol. 195, 3947–3956 (2013). doi:10.1128/JB.00296-13

38. Chatterjee A, Johnson CN, Luong P, Hullahalli K, McBride SW, Schubert AM, Palmer KL, Carlson PE, Duerkop BA. Bacteriophage resistance alters antibiotic-ediated intestinal expansion of enterococci. Infect Immun. 87, e00085–00019 (2019). doi:10.1128/IAI.00085-19

39. Furlan S, Matos RC, Kennedy SP, Doublet B, Serror P, Rigottier-Gois L. Fitness restoration of a genetically tractable *Enterococcus faecalis* V583 derivative to study decoration-related phenotypes of the enterococcal polysaccharide antigen. mSphere. 4, (2019). doi:10.1128/mSphere.00310-19

40. Manson JM, Hancock LE, Gilmore MS. Mechanism of chromosomal transfer of Enterococcus faecalis pathogenicity island, capsule, antimicrobial resistance, and other traits. Proc Natl Acad Sci U S A. 107, 12269–12274 (2010). doi:10.1073/pnas.1000139107

41. Dunny GM. Enterococcal sex pheromones: signaling, social behavior, and evolution. Annu Rev Genet. 47, 457–482 (2013). doi:10.1146/annurev-genet-111212-133449

42. Paulsen IT, Banerjei L, Myers GSA, Nelson KE, Seshadri R, Read TD, Fouts DE, Eisen JA, Gill SR, Heidelberg JF, Tettelin H, Dodson RJ, Umayam L, Brinkac L, Beanan M, Daugherty S, DeBoy RT, Durkin S, Kolonay J, Madupu R, Nelson W, Vamathevan J, Tran B, Upton J, Hansen T, Shetty J, Khouri H, Utterback T, Radune D, Ketchum KA, Dougherty BA, Fraser CM. Role of mobile DNA in the evolution of vancomycin-resistant Enterococcus faecalis. Science. 299, 2071–2074 (2003). doi:10.1126/science.1080613

43. Jeltsch A, Pingoud A. Horizontal gene transfer contributes to the wide distribution and evolution of type II restriction-modification systems. J Mol Evol. 42, 91–96 (1996). doi:10.1007/BF02198833

44. Harms A, Brodersen DE, Mitarai N, Gerdes K. Toxins, targets, and triggers: An overview of toxin-antitoxin biology. Mol Cell. 70, 768–784 (2018). doi:10.1016/j.molcel.2018.01.003

45. Rocha EPC, Bikard D. Microbial defenses against mobile genetic elements and viruses: Who defends whom from what? PLoS Biol. 20, e3001514 (2022). doi:10.1371/journal.pbio.3001514

46. Altschul SF, Gish W, Miller W, Myers EW, Lipman DJ. Basic local alignment search tool. J Mol Biol. 215, 403–410 (1990). doi:10.1016/S0022-2836(05)80360-2

47. Fernandez-Garcia L, Wood TK. Phage-defense systems are unlikely to cause cell suicide. Viruses. 15, (2023). doi:10.3390/v15091795

48. Lopatina A, Tal N, Sorek R. Abortive infection: Bacterial suicide as an antiviral immune strategy. Annu Rev Virol. 7, 371–384 (2020). doi:10.1146/annurev-virology-011620-040628

49. Aframian N, Eldar A. Abortive infection antiphage defense systems: separating mechanism and phenotype. Trends Microbiol. 31, 1003–1012 (2023). doi:10.1016/j.tim.2023.05.002

50. Afonina I, Ong J, Chua J, Lu T, Kline KA. Multiplex CRISPRi system enables the study of stage-specific biofilm genetic requirements in *Enterococcus faecalis*. mBio. 11, e01101–01120 (2020). doi:10.1128/mBio.01101-20

51. Machnicka MA, Kaminska KH, Dunin-Horkawicz S, Bujnicki JM. Phylogenomics and sequence-structure-function relationships in the GmrSD family of Type IV restriction enzymes. BMC Bioinformatics. 16, 336 (2015). doi:10.1186/s12859-015-0773-z

52. Jablonska J, Matelska D, Steczkiewicz K, Ginalski K. Systematic classification of the His-Me finger superfamily. Nucleic Acids Res. 45, 11479–11494 (2017). doi:10.1093/nar/gkx924

53. Bair CL, Black LW. A type IV modification dependent restriction nuclease that targets glucosylated hydroxymethyl ytosine modified DNAs. J Mol Biol. 366, 768–778 (2007). doi:10.1016/j.jmb.2006.11.051

54. He X, Hull V, Thomas JA, Fu X, Gidwani S, Gupta YK, Black LW, Xu SY. Expression and purification of a single-chain Type IV restriction enzyme Eco94GmrSD and determination of its substrate preference. Sci Rep. 5, 9747 (2015). doi:10.1038/srep09747

55. Picton DM, Luyten YA, Morgan RD, Nelson A, Smith DL, Dryden DTF, Hinton JCD, Blower TR. The phage defence island of a multidrug resistant plasmid uses both BREX and type IV restriction for complementary protection from viruses. Nucleic Acids Res. 49, 11257–11273 (2021). doi:10.1093/nar/gkab906

56. Stern A, Sorek R. The phage-host arms race: shaping the evolution of microbes. Bioessays. 33, 43–51 (2011). doi:10.1002/bies.201000071

57. Rocha EP, Danchin A, Viari A. Evolutionary role of restriction/modification systems as revealed by comparative genome analysis. Genome Res. 11, 946–958 (2001). doi:10.1101/gr.gr-1531rr

58. Varadi M, Anyango S, Deshpande M, Nair S, Natassia C, Yordanova G, Yuan D, Stroe O, Wood G, Laydon A, Žídek A, Green T, Tunyasuvunakool K, Petersen S, Jumper J, Clancy E, Green R, Vora A, Lutfi M, Figurnov M, Cowie A, Hobbs N, Kohli P, Kleywegt G, Birney E, Hassabis D, Velankar S. AlphaFold protein structure database: Massively expanding the structural coverage of protein-sequence space with high-accuracy models. Nucleic Acids Res. 50, D439–D444 (2021). doi:10.1093/nar/gkab1061

59. Jumper J, Evans R, Pritzel A, Green T, Figurnov M, Ronneberger O, Tunyasuvunakool K, Bates R, Žídek A, Potapenko A, Bridgland A, Meyer C, Kohl SAA, Ballard AJ, Cowie A, Romera-Paredes B, Nikolov S, Jain R, Adler J, Back T, Petersen S, Reiman D, Clancy E, Zielinski M, Steinegger M, Pacholska M, Berghammer T, Bodenstein S, Silver D, Vinyals O, Senior AW, Kavukcuoglu K, Kohli P, Hassabis D. Highly accurate protein structure prediction with AlphaFold. Nature. 596, 583–589 (2021). doi:10.1038/s41586-021-03819-2

60. Mirdita M, Schutze K, Moriwaki Y, Heo L, Ovchinnikov S, Steinegger M. ColabFold: making protein folding accessible to all. Nat Methods. 19, 679–682 (2022). doi:10.1038/s41592-022-01488-1

61. Kamitori S, Yoshida H. Structure-function relationship of bacterial SH3 domains. In: SH Domains (ed Kurochkina N). Springer, Cham (2015). doi:10.1007/978-3-319-20098-9_4

62. Roberts GA, Stephanou AS, Kanwar N, Dawson A, Cooper LP, Chen K, Nutley M, Cooper A, Blakely GW, Dryden DTF. Exploring the DNA mimicry of the Ocr protein of phage T7. Nucleic Acids Res. 40, 8129–8143 (2012). doi:10.1093/nar/gks516

63. Walkinshaw MD, Taylor P, Sturrock SS, Atanasiu C, Berge T, Henderson RM, Edwardson JM, Dryden DTF. Structure of Ocr from bacteriophage T7, a protein that mimics B-form DNA. Mol Cell. 9, 187–194 (2002). doi:10.1016/s1097-2765(02)00435-5

64. Derakhshani H, Bernier SP, Marko VA, Surette MG. Completion of draft bacterial genomes by long-read sequencing of synthetic genomic pools. BMC Genomics. 21, 519 (2020). doi:10.1186/s12864-020-06910-6

65. Xu S-Y, Corvaglia AR, Chan S-H, Zheng Y, Linder P. A type IV modification-dependent restriction enzyme SauUSI from *Staphylococcus aureus* subsp. aureus USA300. Nucleic Acids Res. 39, 5597–5610 (2011). doi:10.1093/nar/gkr098

66. Ofir G, Melamed S, Sberro H, Mukamel Z, Silverman S, Yaakov G, Doron S, Sorek R. DISARM is a widespread bacterial defence system with broad anti-phage activities. Nat Microbiol. 3, 90–98 (2018). doi:10.1038/s41564-017-0051-0

67. Sterling AJ, Snelling WJ, Naughton PJ, Ternan NG, Dooley JSG. Competent but complex communication: The phenomena of pheromone-responsive plasmids. PLoS Pathog. 16, e1008310 (2020). doi:10.1371/journal.ppat.1008310

68. Licht TR, Laugesen D, Jensen LB, Jacobsen BL. Transfer of the pheromone-inducible plasmid pCF10 among *Enterococcus faecalis* microorganisms colonizing the intestine of mini-pigs. Appl Environ Microbiol. 68, 187–193 (2002). doi:10.1128/AEM.68.1.187-193.2002

69. Huycke MM, Gilmore MS, Jett BD, Booth JL. Transfer of pheromone-inducible plasmids between *Enterococcus faecalis* in the Syrian hamster gastrointestinal tract. J Infect Dis. 166, 1188–1191 (1992). doi:10.1093/infdis/166.5.1188

70. Rice PA. Serine resolvases. Microbiol Spectr. 3, MDNA3-0045-2014 (2015). doi:10.1128/microbiolspec.MDNA3-0045-2014

71. Deep A, Gu Y, Gao YQ, Ego KM, Herzik MA, Jr., Zhou H, Corbett KD. The SMC-family Wadjet complex protects bacteria from plasmid transformation by recognition and cleavage of closed-circular DNA. Mol Cell. 82, 4145–4159 e4147 (2022). doi:10.1016/j.molcel.2022.09.008

72. Jamet A, Touchon M, Ribeiro-Goncalves B, Carrico JA, Charbit A, Nassif X, Ramirez M, Rocha EPC. A widespread family of polymorphic toxins encoded by temperate phages. BMC Biol. 15, 75 (2017). doi:10.1186/s12915-017-0415-1

73. Millman A, Melamed S, Leavitt A, Doron S, Bernheim A, Hor J, Garb J, Bechon N, Brandis A, Lopatina A, Ofir G, Hochhauser D, Stokar-Avihail A, Tal N, Sharir S, Voichek M, Erez Z, Ferrer JLM, Dar D, Kacen A, Amitai G, Sorek R. An expanded arsenal of immune systems that protect bacteria from phages. Cell Host Microbe. 30, 1556–1569 e1555 (2022). doi:10.1016/j.chom.2022.09.017

74. LeRoux M, Srikant S, Teodoro GIC, Zhang T, Littlehale ML, Doron S, Badiee M, Leung AKL, Sorek R, Laub MT. The DarTG toxin-antitoxin system provides phage defence by ADP-ribosylating viral DNA. Nat Microbiol. 7, 1028–1040 (2022). doi:10.1038/s41564-022-01153-5

75. Thurlow LR, Thomas VC, Hancock LE. Capsular polysaccharide production in *Enterococcus faecalis* and contribution of CpsF to capsule serospecificity. J Bacteriol. 191, 6203–6210 (2009). doi:10.1128/jb.00592-09

76. Lefebre MD, Valvano MA. Construction and evaluation of plasmid vectors optimized for constitutive and regulated gene expression in *Burkholderia cepacia* complex isolates. Appl Environ Microbiol. 68, 5956–5964 (2002). doi:10.1128/AEM.68.12.5956-5964.2002

77. Silverman JM, Agnello DM, Zheng H, Andrews BT, Li M, Catalano CE, Gonen T, Mougous JD. Haemolysin coregulated protein is an exported receptor and chaperone of type VI secretion substrates. Mol Cell. 51, 584–593 (2013). doi:10.1016/j.molcel.2013.07.025

78. Cardona ST, Valvano MA. An expression vector containing a rhamnose-inducible promoter provides tightly regulated gene expression in *Burkholderia cenocepacia*. Plasmid. 54, 219–228 (2005). doi:10.1016/j.plasmid.2005.03.004

79. Darling AE, Mau B, Perna NT. progressiveMauve: multiple genome alignment with gene gain, loss and rearrangement. PLoS One. 5, e11147 (2010). doi:10.1371/journal.pone.0011147

80. Gilchrist CLM, Chooi Y-H. clinker & clustermap.js: automatic generation of gene cluster comparison figures. Bioinformatics. 37, 2473–2475 (2021). doi:10.1093/bioinformatics/btab007

81. Goddard TD, Huang CC, Meng EC, Pettersen EF, Couch GS, Morris JH, Ferrin TE. UCSF ChimeraX: Meeting modern challenges in visualization and analysis. Protein Sci. 27, 14–25 (2018). doi:10.1002/pro.3235

82. Holm L. Using Dali for protein structure comparison. Methods Mol Biol. 2112, 29–42 (2020). doi:10.1007/978-1-0716-0270-6_3

83. Camacho C, Coulouris G, Avagyan V, Ma N, Papadopoulos J, Bealer K, Madden TL. BLAST+: architecture and applications. BMC Bioinformatics. 10, 421 (2009). doi:10.1186/1471-2105-10-421

84. Chaumeil PA, Mussig AJ, Hugenholtz P, Parks DH. GTDB-Tk v2: memory friendly classification with the genome taxonomy database. Bioinformatics. 38, 5315–5316 (2022). doi:10.1093/bioinformatics/btac672

