## Supplementary material for "An enterococcal phage protein broadly inhibits type IV restriction enzymes involved in antiphage defense": Combined Supplementary Figures and Tables

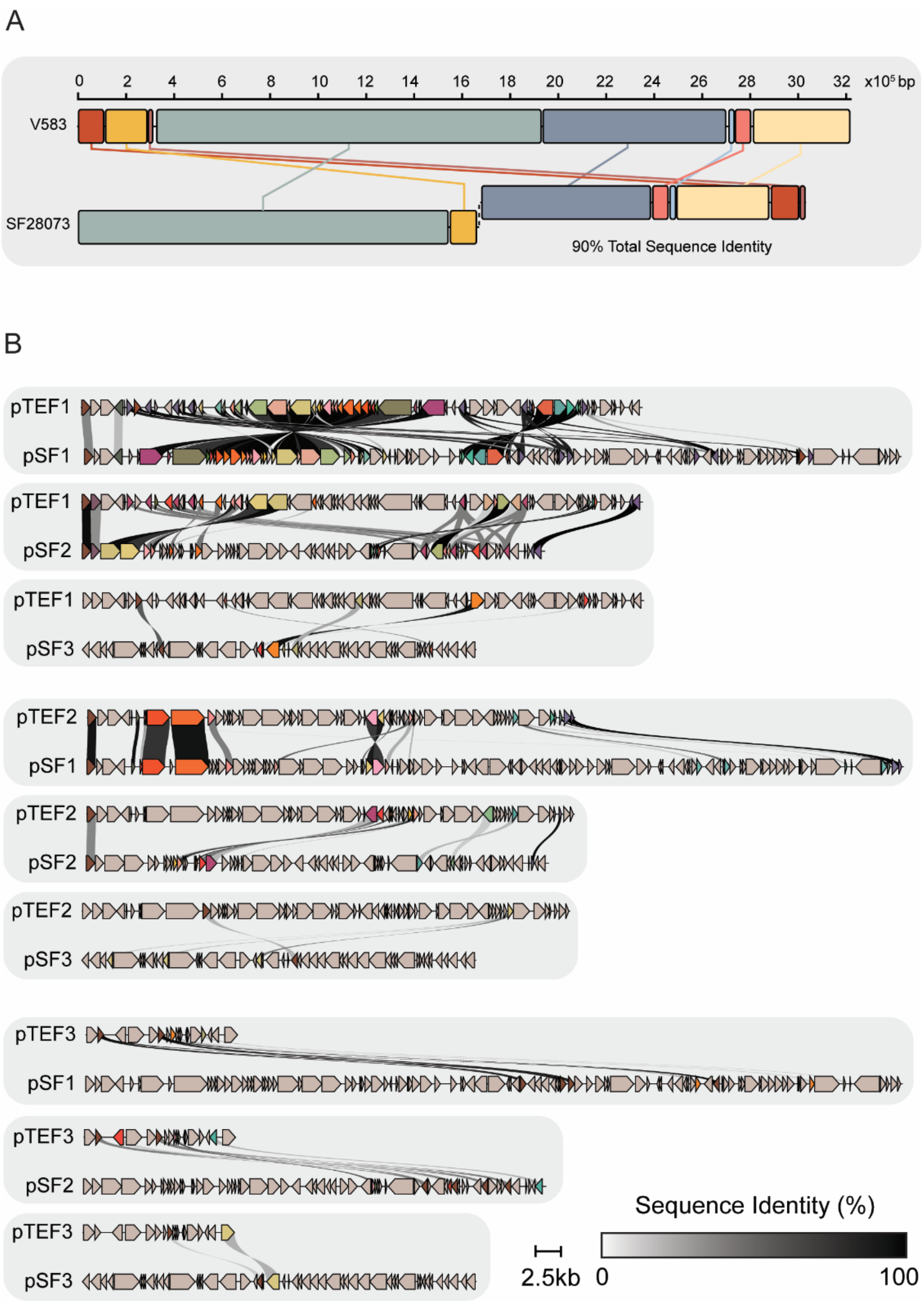


**Figure S1. Genomic comparisons of *E. faecalis* strains V583 and SF28073. (A)** Whole genome alignments of alignments of *E. faecalis* V583 and SF28073 chromosomes **(B)** Sequence alignments between plasmids from *E. faecalis* V583 (pTEF1, pTEF2, pTEF3) and *E. faecalis* SF28073 (pSF1, pSF2, pSF3).

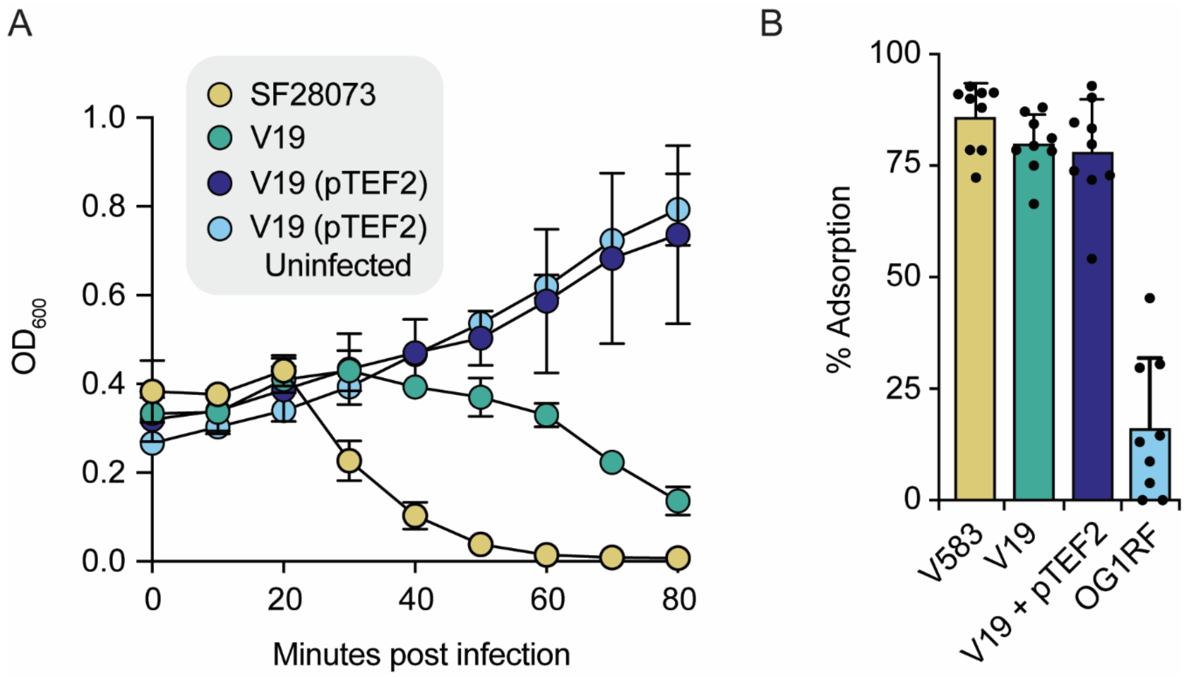


**Figure S2. pTEF2 prevents assembly of intracellular phage particles in an abortive infection-independent manner. (A)** Growth curves of the indicated *E. faecalis* strains infected with Phi47 at an MOI of 10 The growth *E. faecalis* strain V19 containing pTEF2 was measured with and without (uninfected) the addition of phi47. Data represent mean ± s.d. of three technical replicates. (**B)** Percent of phi47 adsorbed to *E. faecalis* strains V583, V19, V19 pTEF2, and OG1RF (negative control). Data represent mean ± s.d. of nine technical replicates. Source data are provided as a Source Data file.

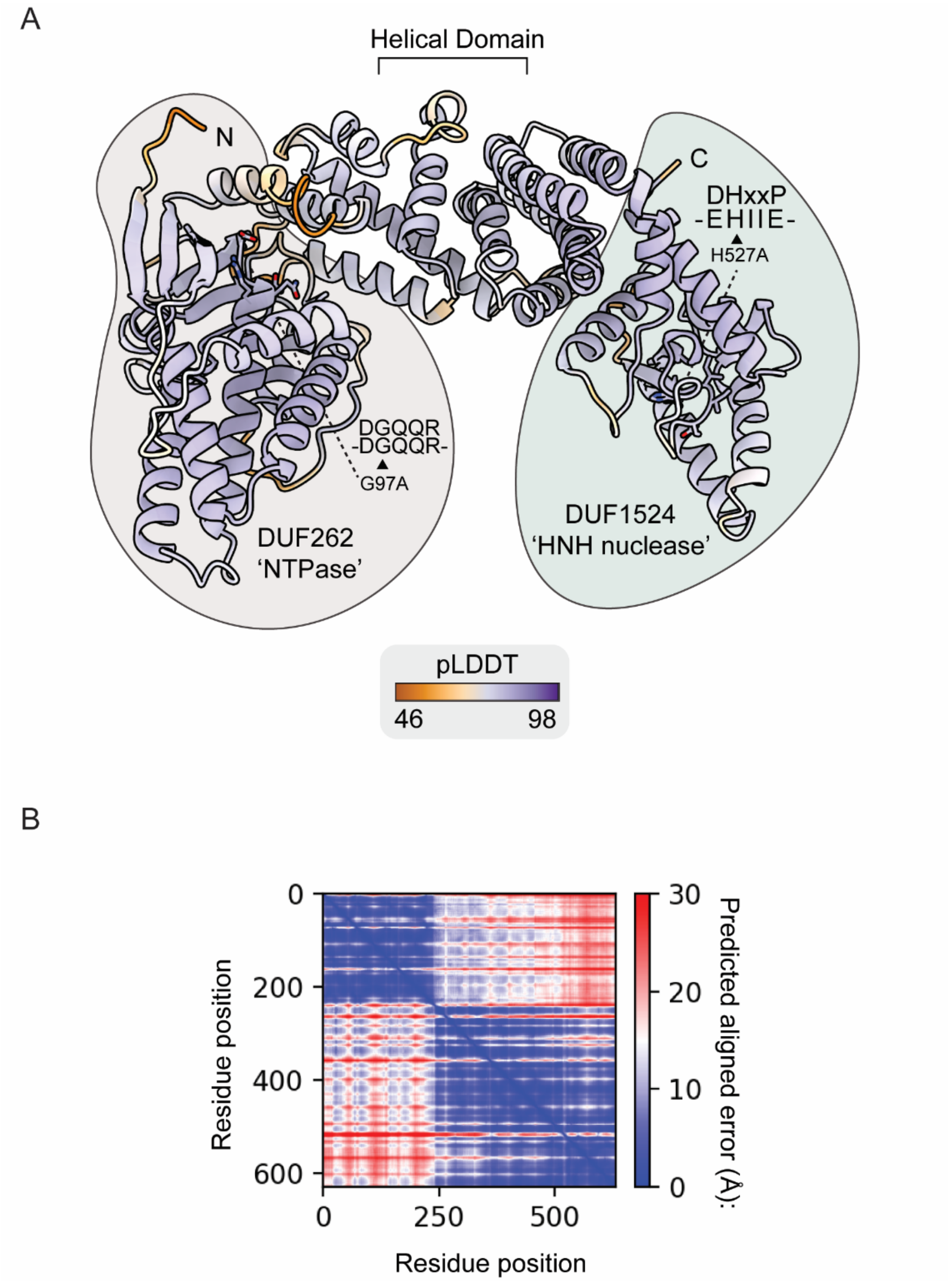


**Figure S3. The predicted structure of EF_B0059 resembles GmrSD-like type IV restriction enzymes. (A)** Ribbon representation of the predicted structure of EF_B0059 generated using Alphafold2. The predicted local distance difference tests (pLDDT) per position is mapped onto the model. Purple corresponds to a more confident prediction at that position. The N-terminal DUF262 and C-terminal DUF1524 domains, and the residues that define each active site are indicated. **(B)** The predicted alignment error (Å) of all residues against all residues for the model in (A). Lower error (blue) corresponds to a well-defined relative position for a given residue pair.

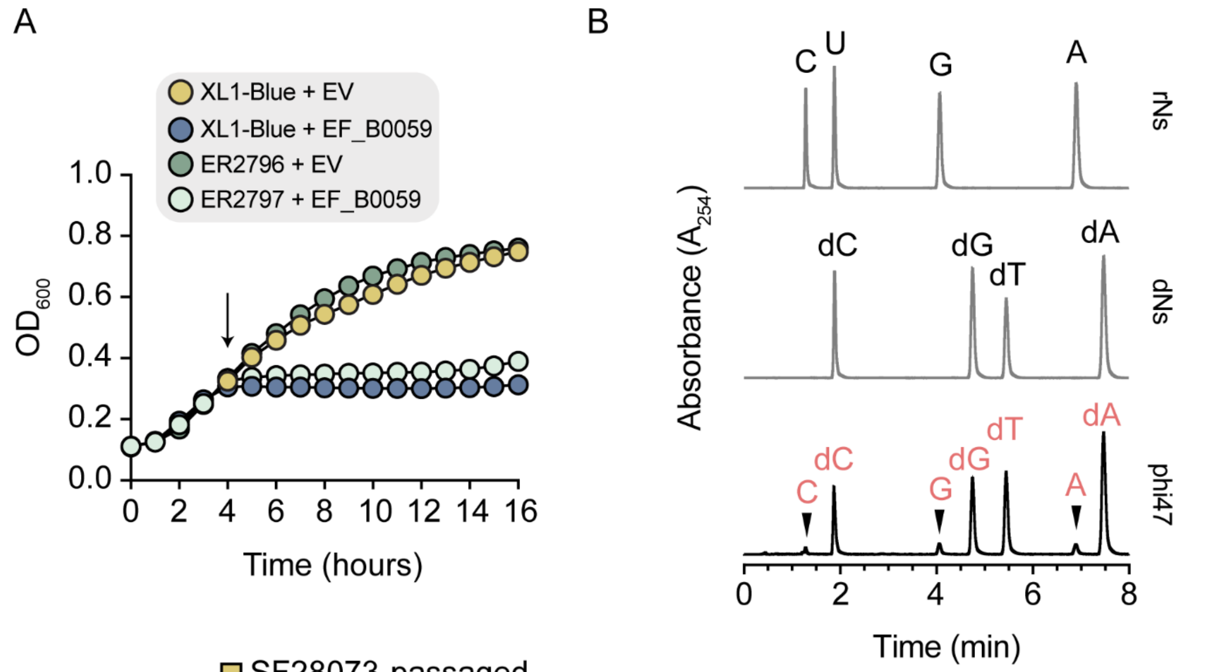


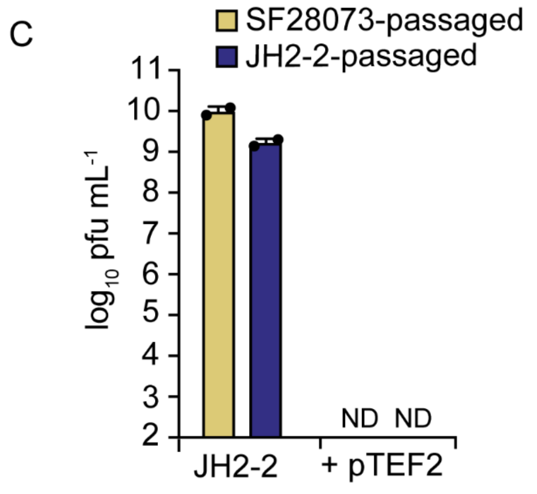


| Product | *m/z* | Measured $[M+H]^+$ | Exact |
|---|---|---|---|
| C | 244.1 | 243.1 | 243.1 |
| dC | 228.1 | 227.1 | 227.1 |
| G | 284.0 | 283.0 | 283.1 |
| dG | 268.0 | 267.0 | 267.1 |
| dT | 243.1 | 242.1 | 242.1 |
| A | 268.1 | 267.1 | 267.1 |
| dA | 252.1 | 251.1 | 251.1 |

**Figure S4. EF_B0059 likely targets unmodified phage DNA. (A)** Growth curves of *E. coli* strains XL1-blue and ER2796 expressing EF_B0059 or empty vector (EV). Arrow indicates when inducer was added to the cultures. Data represent mean ± s.d. of three technical replicates. **(B)** Chromatograms of digested phi47 DNA, deoxyribonucleotide (dNs) standards, or ribonucleotide (rNs) standards. The *m/z*, measured, and expected masses for each peak in the phi47 trace is indicated below **(C)**. Capacity of phage phi47 to form plaques measured by PFU/mL on *E. faecalis* strain JH2-2 or JH2-2 + pTEF2 following passage through the indicated *E. faecalis* strains. Data represent mean ± s.d. of two technical replicates. Source data are provided as a Source Data file.

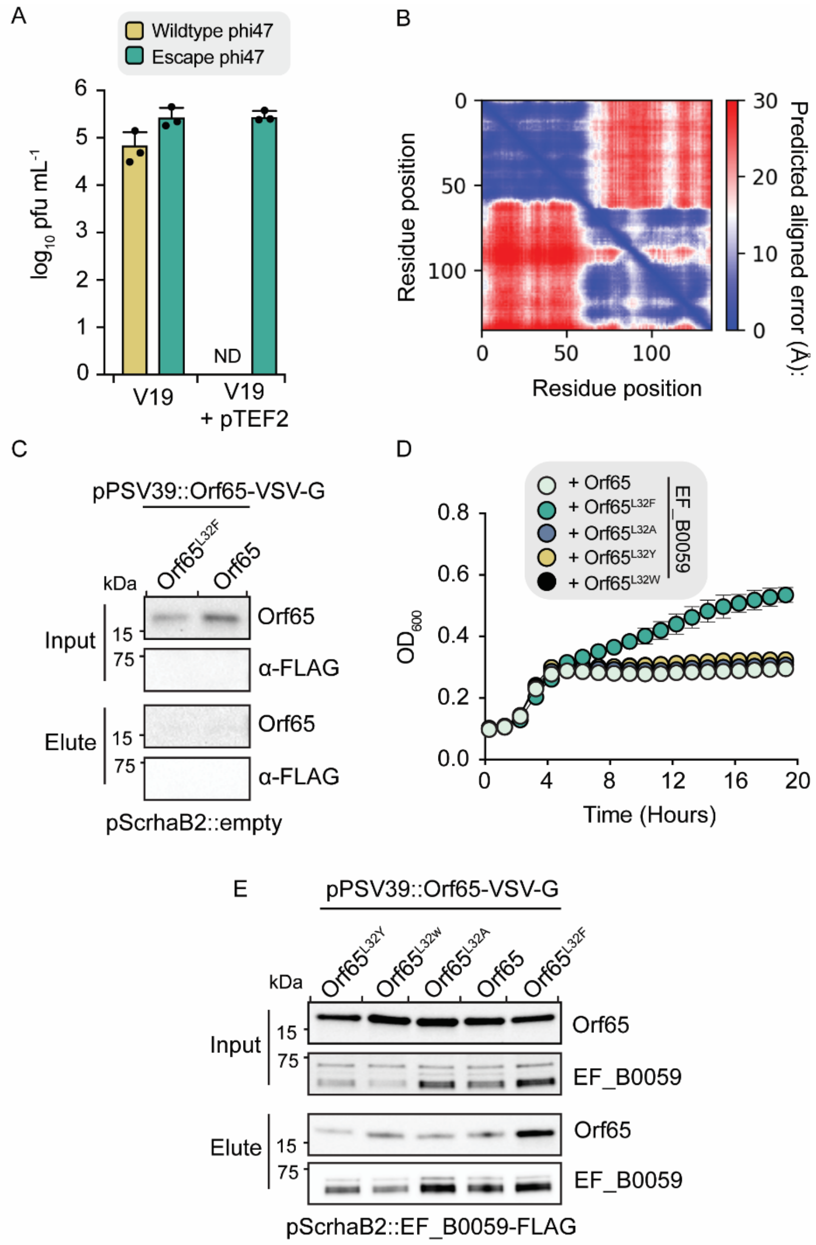


**Figure S5. The L32F mutation in Orf65 is essential for overcoming restriction by EF_B0059. (A)** Outcome of either wildtype of evolved phi47 infections measured by PFU/mL of the *E. faecalis* strain V19 with or without pTEF2. Data represent mean ± s.d. of three technical replicates. **(B)** The predicted alignment error (Å) of all residues against all residues for the structural model in (Fig. 3C). Lower error (blue) corresponds to a well-defined relative position for a given residue pair. **(C)** Anti-FLAG immunoprecipitation of VSV-G tagged Orf65 or Orf65$^{L32F}$ expressed in the absence of EF_B0059-FLAG, detected by Western blot. **(D)** Growth curves of *E. coli* co-expressing EF_B0059 and Orf65 or the indicated Orf65 mutant. Data represent mean ± s.d. of three technical replicates, with the exception of the '+ Orf65$^{L32W}$' sample which represents mean ± s.d. of two technical replicates. **(E)** Anti-FLAG immunoprecipitation of VSV-G tagged Orf65, or the indicated L32 mutant. Source data are provided as a Source Data file.

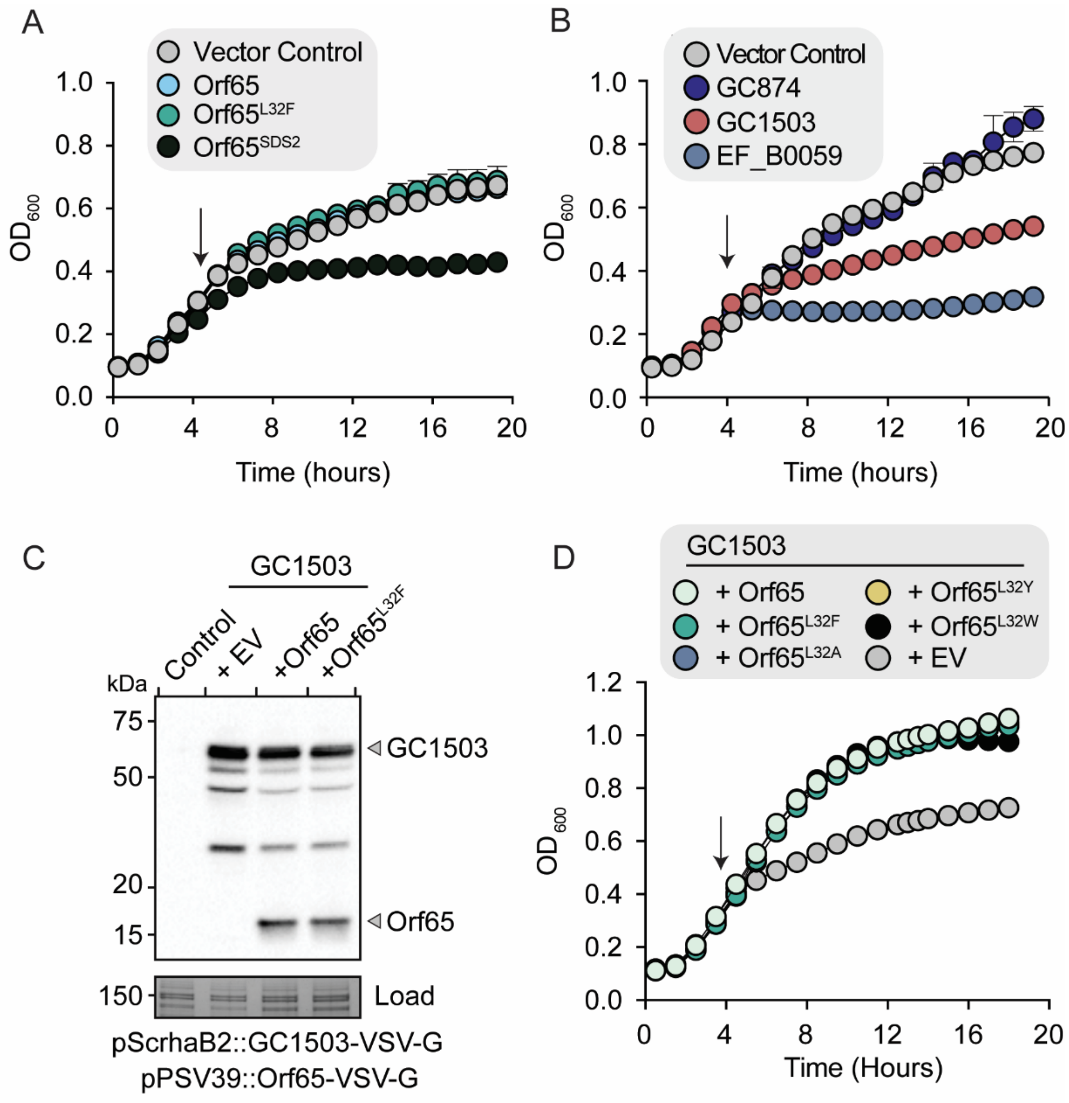


**Figure S6. GC1503, but not GC874, is toxic to *E. coli.* (A)** Growth curves of *E. coli* expressing the indicated Orf65 variants. Orf65$^{SDS2}$ refers to the Orf65 homolog from the enterococcus phage SDS2. Data represent mean ± s.d. of three technical replicates. **(B)** Growth curves of *E. coli* expressing either EF_B0059, GC1503, GC874, or an empty vector control. Arrow indicates when protein expression was induced. Data represent mean ± s.d. of three technical replicates. **(C)** Western blot of VSV-G-tagged EF_B0059 and Orf65 or Orf65$^{L32F}$ in *E. coli* strains 30 minutes post-induction. A non-specific band from the PAGE of these samples stained with Coomassie Brilliant Blue is used as a loading control (load). Data are representative of three independent experiments. **(D)** Growth curves of *E. coli* co-expressing GC1503 and the indicated Orf65 mutant. Arrow indicates when protein expression was induced. Data represent mean ± s.d. of three technical replicates. Source data are provided as a Source Data file.

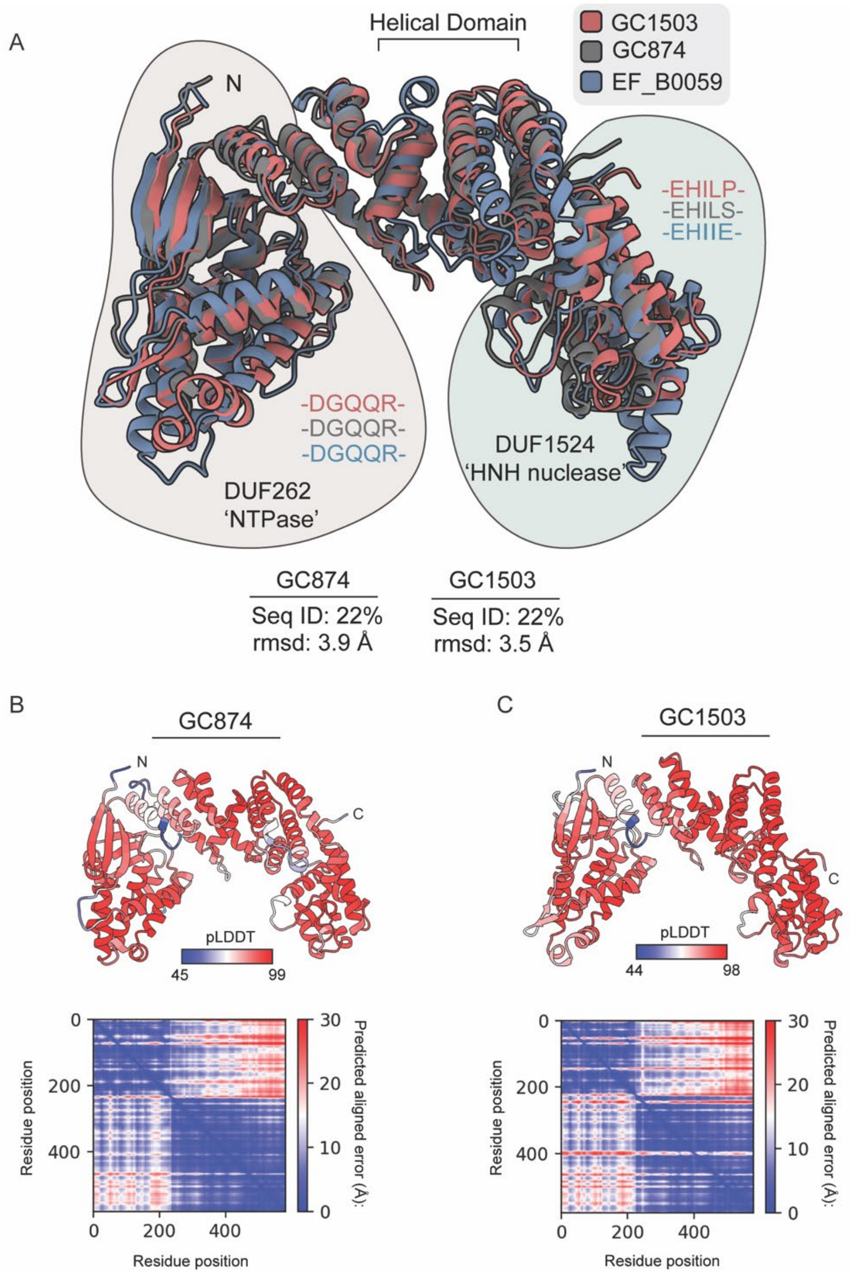


**Figure S7. GC1503 and GC874 are predicted to be structurally similar to EF_B0059. (A)** Structural alignment of the predicted GC1503 (grey) and GC874 (red) models with that of EF_B0059 (blue). Domains corresponding to the N-terminal DUF262 and C-terminal DUF1524 are highlighted, along with the sequences corresponding to the conserved catalytic resides for each structure in these domains. The alignment statistics, including the sequence identify and Cα root mean squared deviations (rmsd) are also indicated. **(B)** Structural model of GC874 (upper) coloured according to the per position predicted local distance difference test (pLDDT) and the predicted alignment error (Å) plot. **(C)** Structural model of GC1503 (upper) coloured according to the per position predicted local distance difference test (pLDDT) and the predicted alignment error (Å) plot.

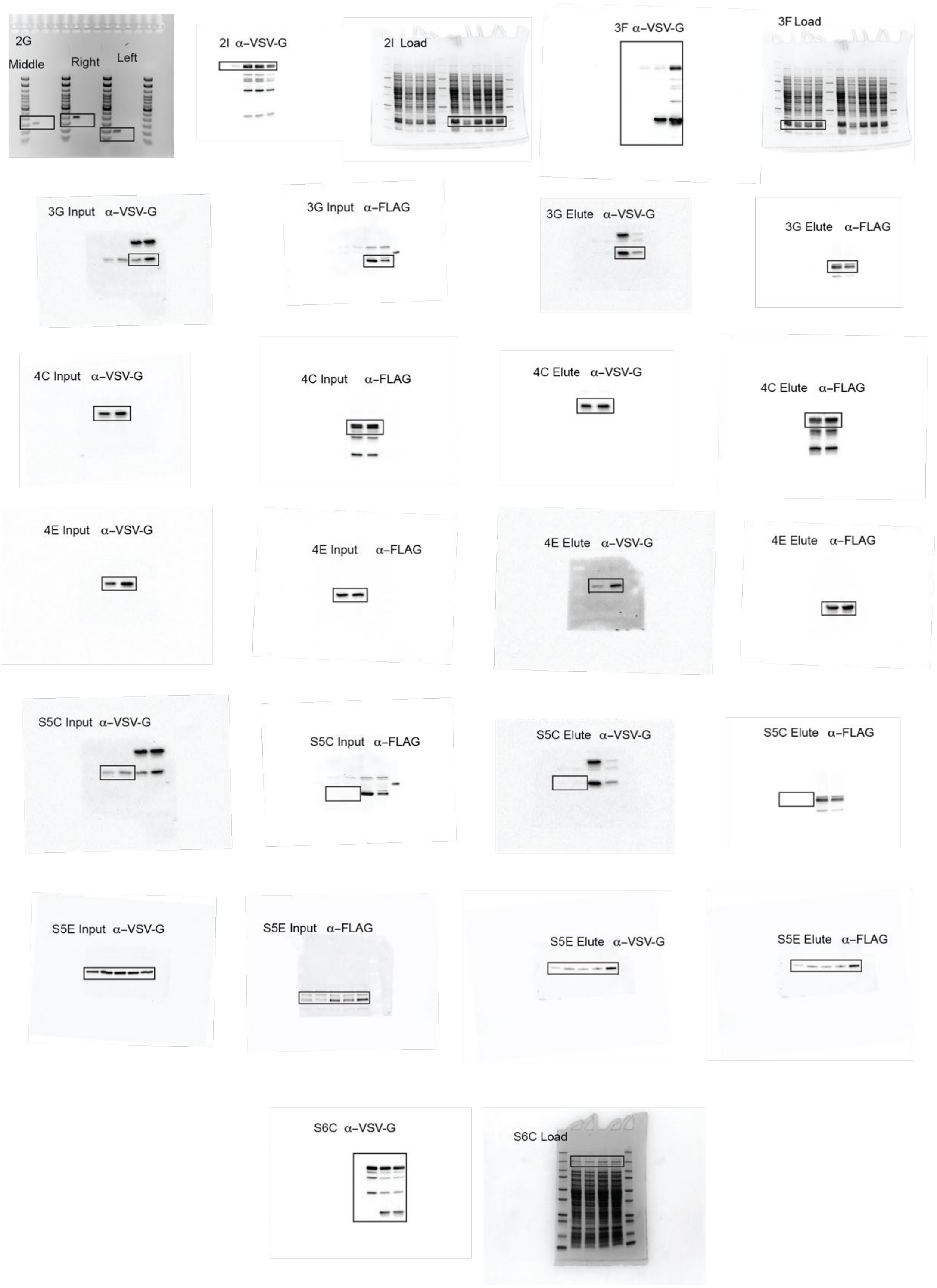


**Figure S8. Full-length, raw versions of gels and blots.** Full-length, unmanipulated versions of all blots and gels presented in the manuscript with corresponding figure numbers. Sections that are displayed in their respective figures are outlined in black. Experiments presented in 2I and 3F, as well as 3G and S5C (control) were performed in tandem and were analyzed using the same gels.

**Table S1. SNPs in Phi47 isolates before and after selection by pTEF2.**

| Isolate ID | Mutation type | Mutation and position | Amino acid change | Coverage | ORF |
|---|---|---|---|---|---|
| Phi47.1 | SNV | G → A, 26201 | Gly812Asp | 99.32 | 33 |
| Phi47.1 pTEF2$^R$ 1 | SNV | G → A, 26201 | Gly812Asp | 99.39 | 33 |
| | **SNV** | **C → T, 46349** | **Leu32Phe** | **99.19** | **65 (TifA)** |
| | Insertion | +GCTAAA, 57289 | - | 36.89 | - |
| Phi47.1 pTEF2$^R$ 2 | SNV | G → A, 26201 | Gly812Asp | 97.80 | 33 |
| | **SNV** | **C → T, 46349** | **Leu32Phe** | **97.35** | **65 (TifA)** |
| Phi47.2 | SNV | G → A, 26201 | Gly812Asp | 99.59 | 33 |
| Phi47.2 pTEF2$^R$ 1 | SNV | G → A, 26201 | Gly812Asp | 99.58 | 33 |
| | **SNV** | **C → T, 46349** | **Leu32Phe** | **99.47** | **65 (TifA)** |
| Phi47.2 pTEF2$^R$ 2 | SNV | G → A, 26201 | Gly812Asp | 99.54 | 33 |
| | **SNV** | **C → T, 46349** | **Leu32Phe** | **99.28** | **65 (TifA)** |
| Phi47.3 | SNV | G → A, 26201 | Gly812Asp | 65.55 | 33 |
| | SNV | T → G, 26342 | Val859Gly | 33.80 | 33 |
| | Deletion | -T, 42638 | Phe33fs | 31.89 | 60 |
| | **SNV** | **C → T, 46349** | **Leu32Phe** | **71.73** | **65 (TifA)** |
| | Insertion | +GCTAAA, 57289 | - | 39.31 | - |
| Phi47.3 pTEF2$^R$ 1 | SNV | T → G, 26342 | Val859Gly | 99.64 | 33 |
| | Deletion | -T, 42638 | Phe33fs | 95.14 | 60 |
| | **SNV** | **C → T, 46349** | **Leu32Phe** | **99.15** | **65 (TifA)** |
| Phi47.3 pTEF2$^R$ 2 | SNV | T → G, 26342 | Val859Gly | 99.72 | 33 |
| | Deletion | -T, 42638 | Phe33fs | 95.63 | 60 |
| | **SNV** | **C → T, 46349** | **Leu32Phe** | **95.65** | **65 (TifA)** |

Three isolates of Phi47 were independently grown to high titer and subjected to selection by pTEF2. Phi47.1, Phi47.2, and Phi47.3 represent the three original stocks of Phi47 prior to selection. pTEF2R 1 and pTEF2R 2 denote each of the two pTEF2-resistant isolates sequenced for each original stock. Orf65 (TifA) mutations are bolded.

**Table S2. Bacterial/bacteriophage strains, plasmids, and oligonucleotides used in this study.**

| Strain/Bacteriophage/plasmid | Relevant properties | Reference or source |
|---|---|---|
| **Bacteria** | | |
| ***E. faecalis*** | | |
| V583 | Human blood isolate; Vanc$^R$, Em$^R$, Gm$^R$ | (1) |
| V19 | Plasmid-cured isolate of *E. faecalis* V583 | (2) |
| JH2-2 | Clinical isolate; Rf$^R$, Fa$^R$ | (3) |
| SF28073 | Human urine sample, 2003, Michigan, USA; Vanc$^R$, Em$^R$, Gm$^R$ | (4) |
| OG1RF | Human oral isolate; Rf$^R$, Fa$^R$ | (5) |
| OG1RF pTEF2-spec$^R$ | Rf$^R$, Fa$^R$, Sp$^R$ | (6) |
| OG1SSp pTEF1 | St$^R$, Sp$^R$, Gm$^R$ | (6) |
| V19 pTEF1 | V19 carrying pTEF1 marked with Sp$^R$ | This study |
| V19 pTEF2 | V19 carrying pTEF2 marked with Sp$^R$ | This study |
| V19 pTEF1, pTEF2 | V19 carrying pTEF1 and pTEF2 marked with Sp$^R$ | This study |
| JH2-2 pTEF2 | JH2-2 carrying pTEF2 marked with Sp$^R$ | This study |
| V19 pCF10 | V19 carrying pCF10, Tc$^R$ | This study |
| V19 pTEF2, pMSP3545-dCas9str | V19 carrying pTEF2 and pMSP3545::dCas9str, Sp$^R$, Em$^R$ | This study |
| V19 pTEF2, pMSP3545-dCas9str, pGCP123* | V19 carrying pTEF2, pMSP3545::dCas9str, and pCGP123 with a spacer for CRISPRi, Sp$^R$, Em$^R$, and Km$^R$ | This study |
| V583 ΔEF_B0058 | V583 *EF_B0058* deletion mutant | This study |
| V583 ΔEF_B0059 | V583 *EF_B0059* deletion mutant | This study |
| V19 pTEF2ΔEF_B0058 | V19 carrying pTEF2 with *EF_B0058* deletion mutant, Sp$^R$ | This study |
| V19 pTEF2ΔEF_B0059 | V19 carrying pTEF2 with *EF_B0059* deletion mutant, Sp$^R$ | This study |
| V19 pTEF2, pLZ12A::orf65 | V19 carrying pTEF2 and pLZ12A expressing phi47 *orf65*, spec$^R$, Cm$^R$ | This study |
| V19 pTEF2, pLZ12A::orf65-L32F | V19 carrying pTEF2 and pLZ12A expressing phi47 *orf65* L32F, Sp$^R$, Cm$^R$ | This study |
| ***E. coli*** | | |
| XL-1 Blue | *recA1 endA1 gyrA96 thi-1 hsdR17 supE44 relA1* lac [F´ *proAB lacI*$^q$ Z$\Delta$*M15* Tn10 (Tet$^R$)] | Novagen |
| ***E. fergusonii*** | | |
| ATCC 35469 | Type strain obtained from ATCC | (7) |
| ***C. sporogenes*** | | |
| GC1503 | Human gut isolate | (8) |
| ***L. mesenteroides*** | | |
| GC874 | Human gut isolate | (8) |
| **Bacteriophages** | | |
| phi47 | Obtained from Washington, DC wastewater | (9) |
| phi47 L32F | pTEF2 escape phi47 mutant, L32F mutation in *orf65* | This study |
| JCW01 | Coliphage isolated from wastewater | This study |

| | | |
|---|---|---|
| SDS2 | Obtained from California wastewater samples | (10) |
| **Plasmids** | | |
| pLT06 | *E. faecalis* allelic exchange vector; $Cm^R$ | (11) |
| pGCP123 | Small shuttle vector for sgRNA expression, $Km^R$ | (12, 13) |
| pMSP3545-dCas9str | Encodes a dead Cas9 from *Streptococcus pyogenes* under the control of nisin-inducible promoter *nisA*, $Em^R$ | (13) |
| pGCP123::GFP | pGCP123 harboring sgRNA for GFP (negative control) | This study |
| pLZ12A | P-*bacA* promoter cloned into shuttle vector pLZ12; pSH71 origin; $Cm^R$ | (9) |
| pGCP123::EF_B0048 | pGCP123 harboring sgRNA for *E. faecalis* V583 gene *EF_B0048* | This study |
| pGCP123::EF_B0049 | pGCP123 harboring sgRNA for *E. faecalis* V583 gene *EF_B0049* | This study |
| pGCP123::EF_B0050 | pGCP123 harboring sgRNA for *E. faecalis* V583 gene *EF_B0050* | This study |
| pGCP123::EF_B0051 | pGCP123 harboring sgRNA for *E. faecalis* V583 gene *EF_B0051* | This study |
| pGCP123::EF_B0052 | pGCP123 harboring sgRNA for *E. faecalis* V583 gene *EF_B0052* | This study |
| pGCP123::EF_B0053 | pGCP123 harboring sgRNA for *E. faecalis* V583 gene *EF_B0053* | This study |
| pGCP123::EF_B0054 | pGCP123 harboring sgRNA for *E. faecalis* V583 gene *EF_B0054* | This study |
| pGCP123::EF_B0055 | pGCP123 harboring sgRNA for *E. faecalis* V583 gene *EF_B0055* | This study |
| pGCP123::EF_B0056 | pGCP123 harboring sgRNA for *E. faecalis* V583 gene *EF_B0056* | This study |
| pGCP123::EF_B0057 | pGCP123 harboring sgRNA for *E. faecalis* V583 gene *EF_B0057* | This study |
| pGCP123::EF_B0058 | pGCP123 harboring sgRNA for *E. faecalis* V583 gene *EF_B0058* | This study |
| pGCP123::EF_B0059 | pGCP123 harboring sgRNA for *E. faecalis* V583 gene *EF_B0059* | This study |
| pGCP123::EF_B0060 | pGCP123 harboring sgRNA for *E. faecalis* V583 gene *EF_B0060* | This study |
| pGCP123::EF_B0061 | pGCP123 harboring sgRNA for *E. faecalis* V583 gene *EF_B0061* | This study |
| pGCP123::EF_B0062 | pGCP123 harboring sgRNA for *E. faecalis* V583 gene *EF_B0062* | This study |
| pGCP123::EF_B0063 | pGCP123 harboring sgRNA for *E. faecalis* V583 gene *EF_B0063* | This study |
| pGCP123::EF_B0064 | pGCP123 harboring sgRNA for *E. faecalis* V583 gene *EF_B0064* | This study |
| pGCP123::EF_B0065 | pGCP123 harboring sgRNA for *E. faecalis* V583 gene *EF_B0065* | This study |
| pLZ12A::*orf65* | pLZ12A constitutively expressing phi47 *orf65*; $Cm^R$ | This study |
| pLZ12A::*orf65*-L32F | pLZ12A constitutively expressing phi47 *orf65* L32F mutant; $Cm^R$ | This study |
| pSCrhaB2-CV | Expression vector with *PrhaB*, C-terminal VSV-G tag, $Tp^R$ | (14) |

| | | |
|---|---|---|
| pSCrhaB2-CV::EF_B0059 | pSCrhaB2-CV harboring EF_B0059 from *E. faecalis* V583 | This study |
| pSCrhaB2-CV::EF_B0059_G97A | pSCrhaB2-CV harboring EF_B0059 G97A mutant | This study |
| pSCrhaB2-CV::EF_B0059_H527A | pSCrhaB2-CV harboring EF_B0059 H527A mutant | This study |
| pSCrhaB2-CV::EF_B0059_G97A_H527A | pSCrhaB2-CV harboring EF_B0059 G97A and H527A mutant | This study |
| pSCrhaB2-CV::EF_B0059-FLAG | pSCrhaB2-CV harboring EF_B0059 with a C-terminal FLAG tag | This study |
| pSCrhaB2-CV::BrxU-FLAG | pSCrhaB2-CV harboring BrxU from *E. fergusonii* ATCC 35469 with a C-terminal FLAG tag | This study |
| pSCrhaB2-CV::GC1503 | pSCrhaB2-CV harboring GC1503 from *C. sporogenes* | This study |
| pSCrhaB2-CV::GC1503-FLAG | pSCrhaB2-CV harboring GC1503 from *C. sporogenes* with a C-terminal FLAG tag | This study |
| pSCrhaB2-CV::GC874 | pSCrhaB2-CV harboring GC1503 from *L. mesenteroides* | This study |
| pPSV39-CV | Expression vector with *lacI, lacUV5* promoter, C-terminal VSV-G tag, $Gm^R$ | (15) |
| pPSV39-CV::orf65 | pPSV39-CV harboring *orf65* from phi47 | This study |
| pPSV39-CV::orf65_L32F | pPSV39-CV harboring *orf65* L32F | This study |
| pPSV39-CV::orf65_L32A | pPSV39-CV harboring *orf65* L32A | This study |
| pPSV39-CV::orf65_L32Y | pPSV39-CV harboring *orf65* L32Y | This study |
| pPSV39-CV::orf65_L32W | pPSV39-CV harboring *orf65* L32W | This study |
| pPSV39-CV::orf65_SDS2 | pPSV39-CV harboring an *orf65* homolog from phage SDS2 | This study |

**Table S3. sgRNA oligonucleotides used in this study**

| Name | Description | Sequence |
|---|---|---|
| EF_B0048 | sgRNA against *E. faecalis* V583 *EF_B0048,* targeting the nontemplate strand, 35% GC | AGCTTAACTAAAATATCTAC |
| EF_B0049 | sgRNA against E. faecalis V583 *EF_B0049*, targeting the nontemplate strand, 30% GC | CAGTTTCCTTTAATTCGTTG |
| EF_B0050 | sgRNA against E. faecalis V583 *EF_B0050*, targeting the nontemplate strand, 30% GC | ATACTATCTTGCATTACATG |
| EF_B0051 | sgRNA against E. faecalis V583 *EF_B0051*, targeting the nontemplate strand, 15% GC | TTACTATTCCATAATAATAT |
| EF_B0053 | sgRNA against E. faecalis V583 *EF_B0053*, targeting the nontemplate strand, 20% GC | AAATAGTTTTTTTGAAGTCT |
| EF_B0054 | sgRNA against E. faecalis V583 *EF_B0054*, targeting the nontemplate strand, 20% GC | TCTTTTTTTTGTACATTACA |
| EF_B0055 | sgRNA against E. faecalis V583 *EF_B0055*, targeting the nontemplate strand, 35% GC | TGGCCTACCAAAATATACAT |
| EF_B0056 | sgRNA against E. faecalis V583 *EF_B0056*, targeting the nontemplate strand, 35% GC | ACAATATCATTACAACCAGC |
| EF_B0057 | sgRNA against E. faecalis V583 *EF_B0057*, targeting the nontemplate strand, 30% GC | ATAATGGATAGATTCCATTC |

| | | |
|---|---|---|
| EF_B0058 | sgRNA against E. faecalis V583 *EF_B0058*, targeting the nontemplate strand, 45% GC | TCAGTTATACCTGTCAACGC |
| EF_B0059 | sgRNA against E. faecalis V583 *EF_B0059*, targeting the nontemplate strand, 40% GC | TTACTATGTGGCCAAACTGT |
| EF_B0061 | sgRNA against E. faecalis V583 EF_B0061, targeting the nontemplate strand, 45% GC | TCACCTAGAGGAAATGCAGT |
| EF_B0064 | sgRNA against E. faecalis V583 *EF_B0064*, targeting the nontemplate strand, 45% GC | TTATCAGTGGCATTTCCCTG |
| EF_B0065 | sgRNA against E. faecalis V583 EF_B0065, targeting the nontemplate strand, 50% GC | TGCACCCAAAAGTCGTTCTG |
| GFP | sgRNA against *gfp,* from Afonina *et al.,* 2020 | CATCTAATTCAACAAGAATT |

### REFERENCES


1. Paulsen IT, Banerjei L, Myers GSA, Nelson KE, Seshadri R, Read TD, Fouts DE, Eisen JA, Gill SR, Heidelberg JF, Tettelin H, Dodson RJ, Umayam L, Brinkac L, Beanan M, Daugherty S, DeBoy RT, Durkin S, Kolonay J, Madupu R, Nelson W, Vamathevan J, Tran B, Upton J, Hansen T, Shetty J, Khouri H, Utterback T, Radune D, Ketchum KA, Dougherty BA, Fraser CM. 2003. Science 299:2071-2074. doi: 10.1126/science.1080613

2. Zhao C, Hartke A, La Sorda M, Posteraro B, Laplace JM, Auffray Y, Sanguinetti M. 2010. Infect Immun 78:3889-97. doi: 10.1128/IAI.00165-10

3. Jacob AE, Hobbs SJ. 1974. J Bacteriol 117:360-72. doi: 10.1128/jb.117.2.360-372.1974

4. Oprea SF, Zaidi N, Donabedian SM, Balasubramaniam M, Hershberger E, Zervos MJ. 2004. J Antimicrob Chemother 53:626-30. doi: 10.1093/jac/dkh138

5. Bourgogne A, Garsin DA, Qin X, Singh KV, Sillanpaa J, Yerrapragada S, Ding Y, Dugan-Rocha S, Buhay C, Shen H, Chen G, Williams G, Muzny D, Maadani A, Fox KA, Gioia J, Chen L, Shang Y, Arias CA, Nallapareddy SR, Zhao M, Prakash VP, Chowdhury S, Jiang H, Gibbs RA, Murray BE, Highlander SK, Weinstock GM. 2008. Genome Biol 9:R110. doi: 10.1186/gb-2008-9-7-r110

6. Manson JM, Hancock LE, Gilmore MS. 2010. Proc Natl Acad Sci U S A 107:12269-74. doi: 10.1073/pnas.1000139107

7. Farmer JJ, 3rd, Fanning GR, Davis BR, O'Hara CM, Riddle C, Hickman-Brenner FW, Asbury MA, Lowery VA, 3rd, Brenner DJ. 1985. J Clin Microbiol 21:77-81. doi: 10.1128/jcm.21.1.77-81.1985

8. Derakhshani H, Bernier SP, Marko VA, Surette MG. 2020. BMC Genomics 21:519. doi: 10.1186/s12864-020-06910-6

9. Chatterjee A, Johnson CN, Luong P, Hullahalli K, McBride SW, Schubert AM, Palmer KL, Carlson PE, Duerkop BA. 2019. Infect Immun 87:e00085-19. doi: 10.1128/IAI.00085-19

10. Wandro S, Ghatbale P, Attai H, Hendrickson C, Samillano C, Suh J, Dunham SJB, Pride DT, Whiteson K. 2022. mSystems 7:e0001922. doi: 10.1128/msystems.00019-22

11. Thurlow LR, Thomas VC, Hancock LE. 2009. J Bacteriol 191:6203-10. doi: 10.1128/jb.00592-09

12. Millman A, Melamed S, Leavitt A, Doron S, Bernheim A, Hor J, Garb J, Bechon N, Brandis A, Lopatina A, Ofir G, Hochhauser D, Stokar-Avihail A, Tal N, Sharir S, Voichek M, Erez Z, Ferrer JLM, Dar D, Kacen A, Amitai G, Sorek R. 2022. Cell Host Microbe 30:1556-1569 e5. doi: 10.1016/j.chom.2022.09.017

13. Afonina I, Ong J, Chua J, Lu T, Kline KA. 2020. mBio 11:e01101-20. doi: 10.1128/mBio.01101-20

14. Cardona ST, Valvano MA. 2005. Plasmid 54:219-28. doi: 10.1016/j.plasmid.2005.03.004

15. Silverman JM, Agnello DM, Zheng H, Andrews BT, Li M, Catalano CE, Gonen T, Mougous JD. 2013. Mol Cell 51:584-93. doi: 10.1016/j.molcel.2013.07.025
